# “DOCK3 is a dosage-sensitive regulator of skeletal muscle and Duchenne muscular dystrophy-associated pathologies”

**DOI:** 10.1101/2020.03.27.010223

**Authors:** Andrea L. Reid, Yimin Wang, Adrienne Samani, Rylie M. Hightower, Michael A. Lopez, Shawn R. Gilbert, Lara Ianov, David K. Crossman, Louis J. Dell’Italia, Douglas P. Millay, Thomas van Groen, Ganesh V. Halade, Matthew S. Alexander

**Affiliations:** Department of Pediatrics, Division of Neurology at the University of Alabama at Birmingham and Children’s of Alabama, Birmingham, AL 35294; UAB Center for Exercise Medicine, Birmingham, AL 35294; Department of Orthopedic Surgery, the University of Alabama at Birmingham, Birmingham, AL, 35294; Civitan International Research Center at the University of Alabama at Birmingham, Birmingham, AL 35294; Department of Genetics at the University of Alabama at Birmingham, Birmingham, AL 35294; Birmingham Veteran Affairs Medical Center, Birmingham, AL 35233; Department of Medicine, Division of Cardiovascular Disease, the University of Alabama at Birmingham, Birmingham, AL 35294; Department of Cell, Developmental, and Integrative Biology, the University of Alabama at Birmingham, Birmingham, AL 35294; Division of Molecular Cardiovascular Biology, Cincinnati Children’s Hospital Medical Center, Cincinnati, Ohio, 45229; Department of Medicine, Division of Cardiovascular Sciences, the University of South Florida, Tampa, FL, 33602

**Keywords:** DOCK3, myogenic differentiation, DMD, skeletal muscle, dystrophy

## Abstract

DOCK3 is a member of the DOCK family of guanine nucleotide exchange factors that function to regulate cell migration, fusion, and overall viability. Previously, we identified a miR-486/Dock3 signaling cascade that was dysregulated in dystrophin-deficient muscle which resulted in the overexpression of *DOCK3*, however not much else is known about the role of DOCK3 in muscle. In this work, we characterize the functional role of DOCK3 in normal and dystrophic skeletal muscle. By utilizing *Dock3* global knockout (*Dock3* KO) mice, we found reducing *Dock3* gene via haploinsufficiency in DMD mice improved dystrophic muscle histology, however complete loss of *Dock3* worsened overall muscle function on a dystrophin-deficient background. Consistent with this, *Dock3* KO mice have impaired muscle architecture and myogenic differentiation defects. Moreover, transcriptomic analyses of *Dock3* knockout muscles reveal a decrease in factors known for myogenesis, suggesting a possible mechanism of action. These studies identify *DOCK3* as a novel modulator of muscle fusion and muscle health and may yield additional therapeutic targets for treating dystrophic muscle symptoms.

## Introduction

Skeletal muscle formation requires a complex coordination of various signaling factors and cell populations to efficiently form mature myofibers capable of generating muscle force. Duchenne muscular dystrophy (DMD) is an X-linked neuromuscular disorder caused by pathogenic mutations in the *DYSTROPHIN* gene resulting in the lack of functional dystrophin protein. This dystrophin-deficiency results in a degradation of the link between filamentous actin and the extracellular matrix (ECM) causing structural damage. In DMD patients, after the loss of ambulation their skeletal muscle begins to progressively deteriorate and is replaced with adipocytes, inflammatory cells, and necrotic tissue. While the dystrophin protein has been thought of as a classical muscle structural protein, recent evidence suggests that it also plays a significant role in the regulation of various cellular signaling pathways involved in dystrophic pathology^1-5^.

The DOCK (dedicator of cytokinesis) family of proteins function as key signaling factors that regulate cellular migration, fusion, and survival in different cell and tissue types^6^. DOCK1 and DOCK5 have been shown to play key roles in myoblast fusion through Crk1/Mapk14 protein interactions^7,8^. DOCK3 (previously called MOCA or modifier of cell adhesion) was shown to be essential for normal neuronal function and *Dock3* global knockout mice develop plaques in the brain^9^. Human patients with loss-of-function *DOCK3* variants develop intellectual disability, ataxia, developmental delay, and have overall low muscle tone from birth^10-12^. There is strong genetic evidence that the disease severity of patients with *DOCK3* pathogenic variants is linked to the degree of which DOCK3 can activate RAC1 signaling as a guanine nucleotide exchange factor (GEF) protein^13^. Previously, we identified a muscle-enriched microRNA, miR-486, as a regulator of dystrophin-deficient skeletal muscle pathology in mice and human muscle biopsies through its negative regulation of *DOCK3* mRNA expression levels^14,15^. We hypothesized that DOCK3 may be a direct regulator of muscle function and that its dysregulation in DMD may play significant downstream consequences in the progression of the disease. We assessed the roles of DOCK3 in DMD pathology in both zebrafish and mouse models of the disease. We analyzed *Dock3* KO mouse muscles and found significant perturbations in overall muscle integrity and function. Furthermore, *Dock3* KO mouse muscle cells were analyzed for their ability to proliferate, differentiate, and overall viability in culture. Transcriptomic analyses of *Dock3* KO *tibialis anterior* (TA) muscles revealed a significant downregulation of several muscle pathways including the fusogen myomixer (*Mymx*), also referred to as myomerger or minion^16-18^. This work demonstrates the critical role of DOCK3 in normal myogenesis and a gene-dosing effect of DOCK3 expression levels on dystrophin-deficient muscle health.

## Results

### Dock3 gene expression is upregulated in dystrophic muscles

We first aimed to assess endogenous *Dock3* gene expression levels in normal and dystrophin-deficient muscle. Previously, we established *DOCK3* as a mRNA target of miR-486, a muscle-enriched miRNA whose expression is reduced in with dystrophic disease progression^14^. We evaluated *Dock3* gene expression in the DMD mouse model, *mdx*^*5cv*^, and compared to wild type (*C57BL/6J* strain) aged-matched male mice. The *mdx*^*5cv*^ mice have significantly fewer revertant dystrophic fibers, increased muscle weakness, and a reported overall greater dystrophic pathology than the classical *mdx* strain of mice^19,20^. We observed *Dock3* mRNA expression to be significantly elevated in the *mdx*^*5cv*^ muscles compared to aged-matched wild type control muscles across three different muscles types including the TA, soleus, and diaphragm starting at 1 month of life (**Figures 1A-C**). These findings of elevated Dock3 mRNA were found to be consistent throughout adulthood into aged (12-month-old) *mdx*^*5cv*^ muscles (**Figures 1A-C**).

**Figure 1.**
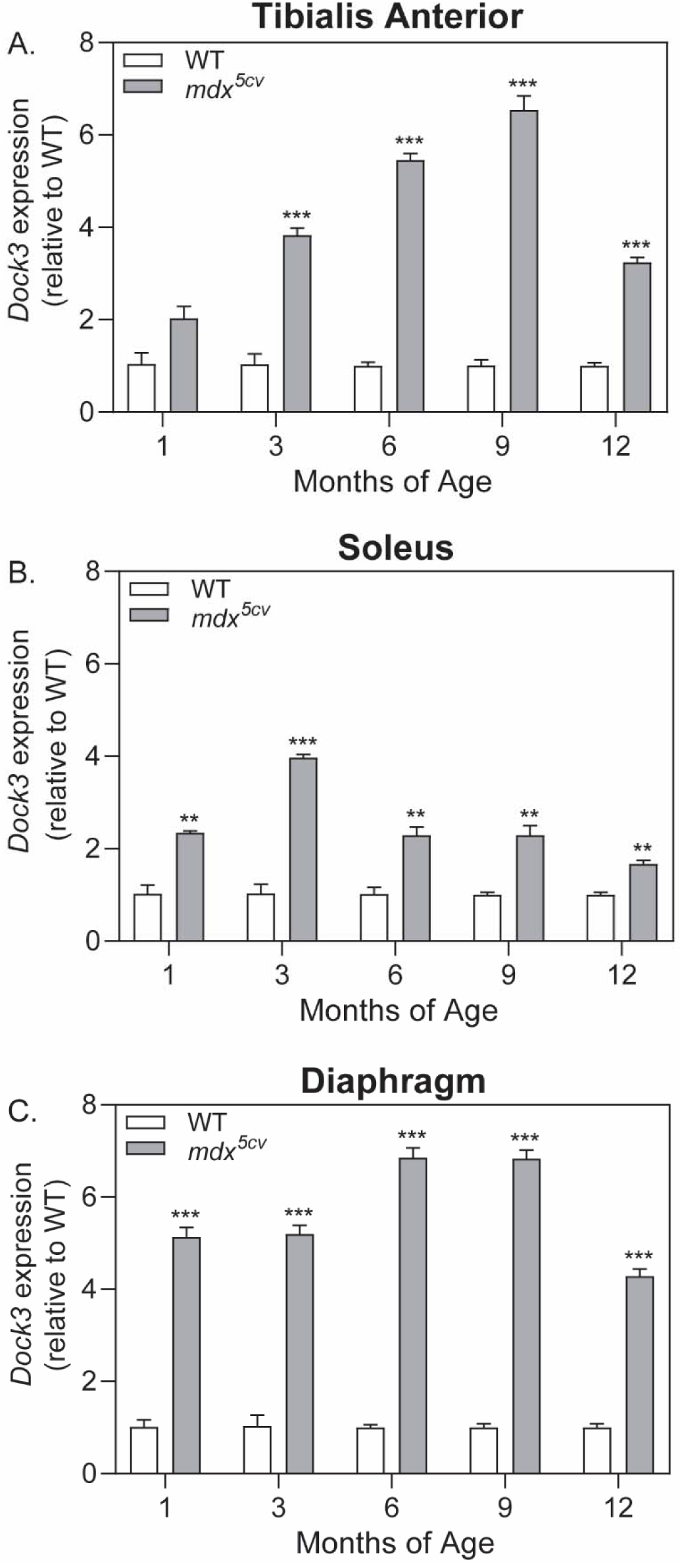
*Dock3* mRNA expression levels are increased in adult *mdx*^*5cv*^ mouse skeletal muscles. Real time qPCR data from wild type (WT) and *mdx*^*5cv*^ mice taken from **A.** *Tibialis anterior* (TA), **B.** soleus, and **C.** diaphragm muscles at 1, 3, 6, 9, and 12 months of age demonstrating the significant increase in *Dock3* mRNA expression in *mdx*^*5cv*^ dystrophic mouse muscle compared to WT muscle. (n=3 per genotype and cohort, performed in triplicate; two-way ANOVA with Tukey’s correction; **p<0.01, ***p<0.001).

### Knockdown of dock3 mRNA in dystrophic zebrafish larvae improves muscle and overall outcomes

Zebrafish are a powerful model system for DMD disease modeling due to their translucent *ex vivo* development, large offspring sizes, ease of genetic manipulation, and ability to uptake compounds in their skin and gills during early developmental stages^21-23^. Zebrafish Dock3 protein amino acid sequences showed a strong amount of evolutionary conservation to mouse and human DOCK3 protein sequences (85% homology with mouse and human; **Supplemental Figure S1**). We performed *in situ* hybridization (ISH) for zebrafish *dock3* mRNA expression in wild type (AB strain) zebrafish embryos at the early stages of development including prior to and after muscle formation. Zebrafish *dock3* mRNA was detected at low levels during early somite formation with strong expression in the developing brain, neural tube, and dorsal muscle by 4 days post fertilization (dpf) (**Figure 2A**). The *sapje* zebrafish is a widely used DMD model that has the severe muscle weakness and myofiber degradation comparable to those observed in the later stages of DMD patient skeletal muscles^21,24,25^. Unlike mammals, zebrafish lack definitive sex chromosomes and the *dystrophin* (*dmd*) gene in zebrafish is located on chromosome 1^26^. Mating of *sapje* heterozygote fish results in a Mendelian percentage of twenty-five percent affected zebrafish that can be easily identified via birefringence assay^24,25^. The *sapje* homozygote mutant larvae showed severe myofiber disruption in affected larvae versus unaffected (*sapje* heterozygotes and wild type) larvae^25,27-29^ (**Figure 2B**). Given our findings of increased *Dock3* expression in *mdx*^*5cv*^ muscles, we tested whether modulation of zebrafish *dock3* expression might affect overall dystrophic outcomes. We observed significantly fewer than expected affected *sapje* zebrafish in our *dock3* morphant (*dock3* MO) zebrafish compared to control morphants when using a low dose of *dock3* morpholino (1 ng) (**Figure 2C**). However, the high dose of *dock3* morpholino (6 ng) resulted in a higher number of affected *sapje* zebrafish, suggesting there may be an optimal dose for blocking dystrophic pathology. These results were further confirmed via histochemical staining of F-actin in the dorsal muscles of the 1 ng *dock3* MO *sapje* zebrafish which revealed fewer areas of myofiber detachment and overall more integrity compared to control MO *sapje* homozygotes, while the converse was observed for the high dose *dock3* morphant (**Figure 2D**). Together, these data suggest that DOCK3 expression levels are strongly linked towards dystrophic pathologies and overall outcomes in DMD zebrafish.

**Figure 2.**
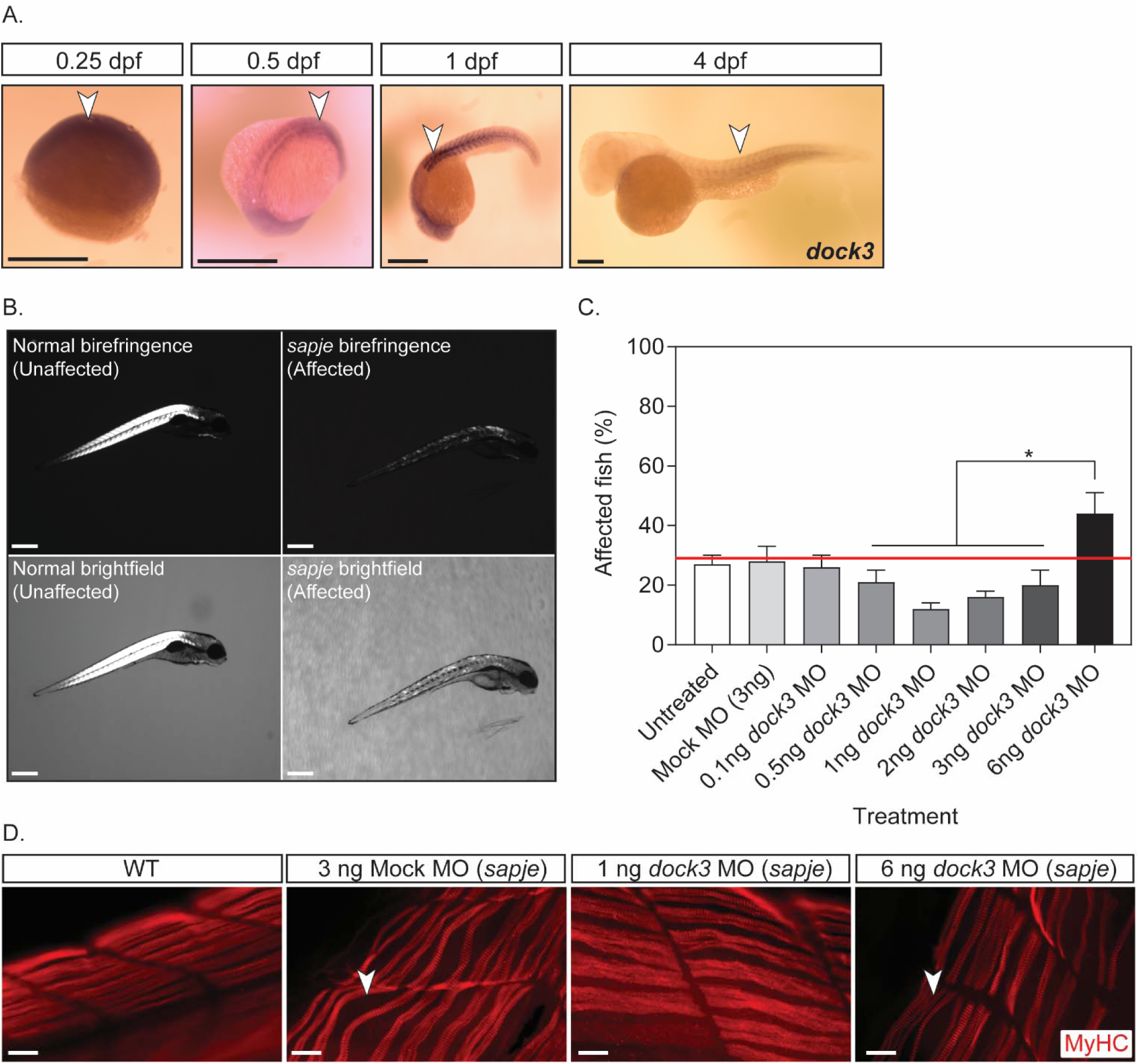
Knockdown of *dock3* mRNA in DMD zebrafish reduces dystrophic pathology. **A.** *In situ hybridization* (ISH) of zebrafish *dock3* mRNA during early developmental time points. Note the robust expression of zebrafish *dock3* mRNA in muscle tissues at stages of early muscle formation and muscle pioneer cell fusion. Not shown sense probes are used as internal controls. Arrowheads demarcate *dock3* mRNA ISH signal. Scale bar = 100 µm. **B.** Representative images of normal and *sapje* mutant birefringence morphology. Scale bar = 200 µm. **C.** Quantification of muscle birefringence shows that 1ng of *dock3* MO improves muscle birefringence scoring in *sapje* mutant zebrafish, while 6ng of *dock3* MO worsens muscle birefringence scoring. (n=3 experimental replicates; one-way ANOVA with Tukey’s correction; *p<0.05). **D.** Myosin heavy chain (MyHC; F-59 antibody) immunofluorescent staining of 4 dpf WT (uninjected) or *sapje* mutant larvae injected with control (mock) morpholino, or *dock3* MO (1 ng or 6 ng). Arrowheads demarcate myofiber tears from the sarcolemmal membrane. Scale bar = 40 µm.

### Knockdown of DOCK3 in isolated DMD myoblasts increases myogenic fusion

To investigate the effects of DOCK3 on myoblast fusion in dystrophin-deficient cells we infected DMD and normal primary human myoblasts with lentiviral particles of either shRNA *DOCK3*, shRNA scrambled (i.e. non-targeting), or a mock control (**Figure 3A**). DMD myoblasts have poor terminal differentiation into myotubes^30,31^. We observed a significantly higher myogenic fusion index in the shRNAi *DOCK3* DMD myotubes compared to either shRNA scrambled or mock-treated DMD muscle cells (**Figures 3A and B**). To validate the *DOCK3* mRNA knock-down, we performed western blot analysis for DOCK3 protein levels and confirmed a marked reduction of DOCK3 protein expression (**Figure 3C**). This data reveals that knockdown of *DOCK3* mRNA in human DMD myoblasts improves myogenic fusion percentages despite dystrophin-deficiency.

**Figure 3.**
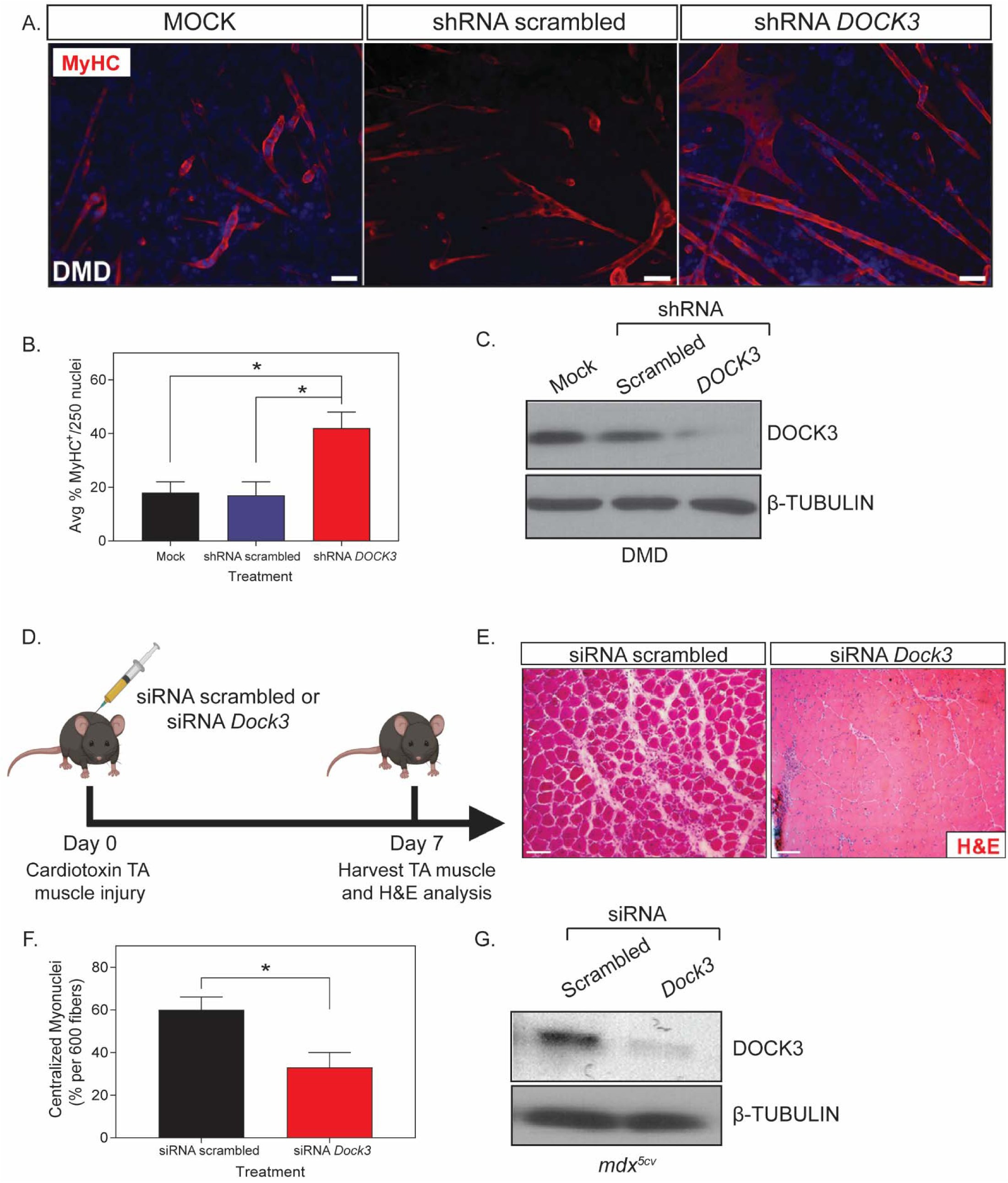
Knockdown of DOCK3 in DMD patient muscle cells and *mdx*^*5cv*^ mice increases myogenic differentiation and improves pathology. Myosin heavy chain staining reveals that knockdown of DOCK3 protein in human DMD primary myotubes increases myogenic fusion indices. **A.** Immunofluorescence of MyHC (MF-20 antibody, DSHB Iowa, red) and nucleus marker (DAPI, blue) of day 7 differentiated human DMD primary myotubes either transduced with mock (control), shRNA scrambled (negative control), and shRNA *DOCK3* lentiviral particles (transduced at a MOI (multiplicity of infection) of 10). Scale bar = 50 µm. **B.** Summary graph of average myogenic fusion indices as calculated by percentage of nuclei inside of MyHC^+^ cells out of 250 nuclei, as previously described^63^. DMD primary myotubes transduced with shRNA *DOCK3* demonstrated significantly higher amount of myogenic fusion. (n=3 experimental replicates, one-way ANOVA with Tukey’s correction, *p<0.05). **C.** Western blot analysis of whole cell lysates taken from day 7 differentiated DMD myotubes transduced with mock (control), shRNA scrambled (shRNA internal control) or shRNA DOCK3 knockdown lentiviral particles, demonstrating successful knockdown of DOCK3 protein expression. **D.** Schematic showing co-injection in *mdx*^*5cv*^ mice of cardiotoxin (10 µM ctx) as well as either siRNA double-stranded (ds) oligos to inhibit mouse *Dock3* mRNA (siRNA *Dock3*) or a scrambled ds oligo (siRNA scrambled; control). Experiment was performed in each leg and mice were evaluated 7 days post ctx injury. **E.** Representative H&E staining of the siRNA-injected *mdx*^*5cv*^ muscles 7 days post ctx. Scale bar = 50 µm. **F.** Summary graph showing reduced centralized myonuclei when *mdx*^*5cv*^ mice were injected with siRNA *Dock3* (n=5, student’s t-test, two-tailed, *p<0.05). **G.** Western blot analysis of whole muscle lysates from siRNA scrambled and siRNA *Dock3* muscle demonstrating successful knockdown of Dock3 protein.

### Knockdown of Dock3 in mdx^5cv^ mice improves cardiotoxin-induced muscle injury repair

We next investigated the role of DOCK3 in DMD pathology using the *mdx*^*5cv*^ mouse model of DMD. Given our findings of DOCK3 in myoblast fusion, we hypothesized DOCK3 might play a role in muscle repair following injury. *Mdx*^*5cv*^ mice were co-injected with cardiotoxin (ctx) into the TA muscles along with either siRNA scrambled (control) or mouse *Dock3* siRNA (**Figure 3D**). The *mdx*^*5cv*^ mice treated with *Dock3* siRNA had a more improved muscle injury repair pathology and muscle architecture compared to siRNA scrambled-treated mice seven days after injury (**Figure 3E**). The siRNA *Dock3-*treated *mdx*^*5cv*^ mice had significantly less centralized myonuclei than the control group (**Figure 3F**). To validate *Dock3* mRNA knockdown, we performed western blot analysis which confirmed a strong reduction of DOCK3 protein expression levels (**Figure 3G**). These results support a potential beneficial effect of reducing DOCK3 expression in the *mdx*^*5cv*^ mice during muscle injury-induced repair and regeneration.

### Haploinsufficiency of Dock3 improves muscle pathologies in mdx^5cv^ mice

Given our previous findings demonstrating that DOCK3 expression levels are elevated in DMD patient and *mdx*^*5cv*^ mouse we used *Dock3* global knockout (KO) mice and evaluated their muscles on wild type and dystrophic *mdx*^*5cv*^ strain backgrounds in six separate cohorts. Muscle architecture was assessed at 6 months of age across all groups by measuring number of centralized myonuclei and fiber size distribution via hematoxylin and eosin (H&E) histochemistry (**Figure 4A**). Global *Dock3* KO mice had increased centralized myonuclei and altered fiber size distribution, indicating impaired muscle health compared to wild type control mice (**Figures 4B, 4C**). In addition, we observed significant muscle fiber group atrophy reflected in smaller myofiber frequencies in the *Dock3* KO mouse muscle compared to control WT muscles (**Figures 4C**). Consistent with these initial findings, the global *Dock3* KO mice on *mdx*^*5cv*^ background [*Dock3:mdx*^*5cv*^ double knockout (DKO)] mice developed exacerbated muscle weakness and all *Dock3:mdx*^*5cv*^ DKO mice expired by six months of age due to diaphragm muscle weakness (data not shown). Muscle structural analyses revealed severe myofiber disruption and fibrosis in the *Dock3:mdx*^*5cv*^ DKO mice compared to *mdx*^*5cv*^ and *Dock3* KO muscles (**Figures 4A, 4D, 4E**). Interestingly, the haploinsufficiency of *Dock3* in *mdx*^*5cv*^ mice (*Dock3*^*+/-*^:*mdx*^*5cv*^) improved muscle fiber architecture, larger myofiber sizes, and overall improved muscle pathology in comparison with the *mdx*^*5cv*^ and *Dock3:mdx*^*5cv*^ DKO double mutant mice (**Figures 4A, 4D, 4E**). These findings suggest that *Dock3* dosage levels influence the overall outcome of normal and dystrophin-deficient muscles, and that haploinsufficiency of *Dock3* is beneficial in dystrophic muscle pathology.

**Figure 4.**
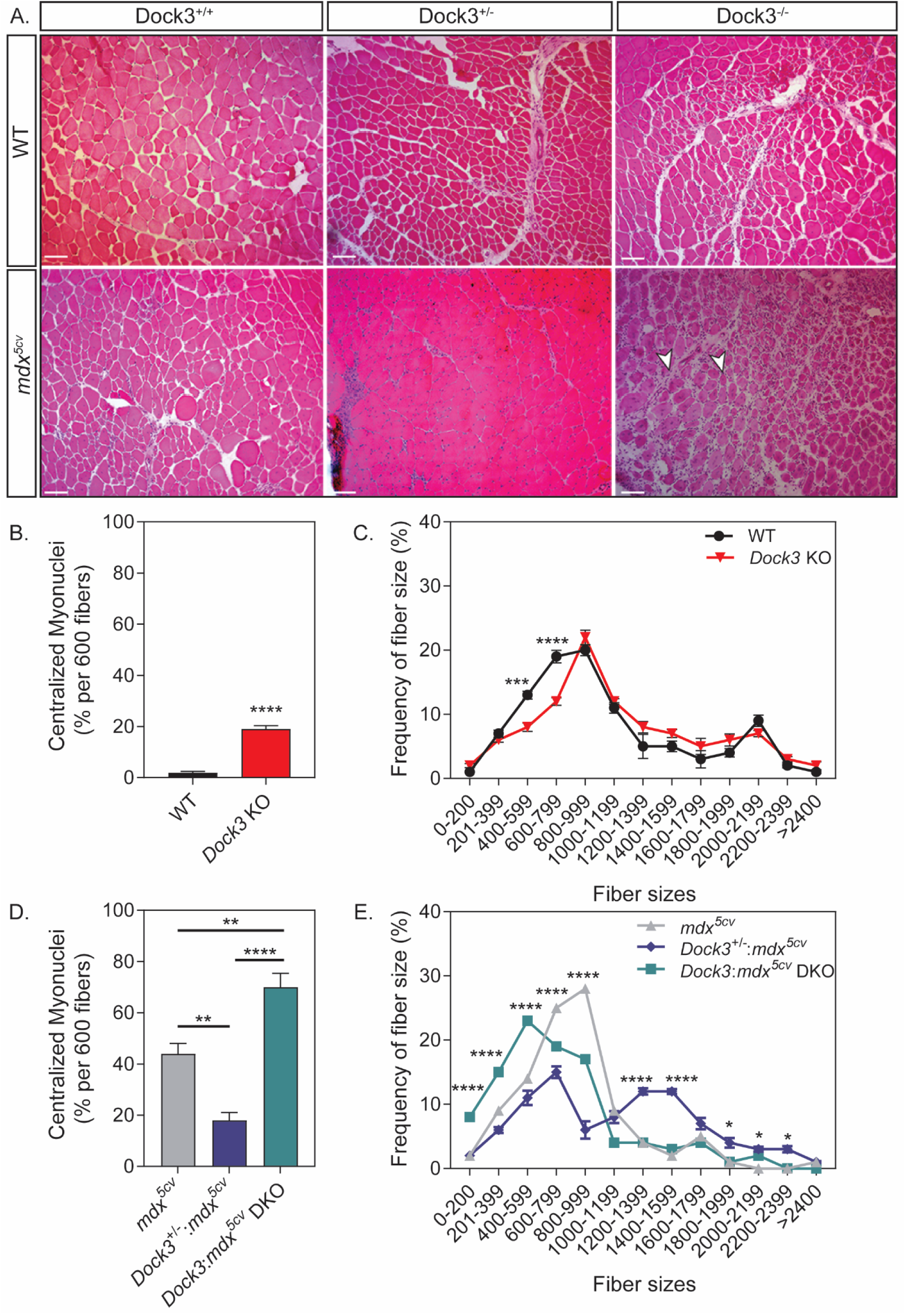
Gene dosage of *Dock3* affects muscle architecture and dystrophic pathology in wild type and *mdx*^*5cv*^ mice. **A.** H&E immunohistochemistry of *Dock3+/+* (wild type), *Dock3+/-*, and *Dock3* KO TA adult muscle on wild type (for dystrophin) and *mdx*^*5cv*^ strain backgrounds. Scale bar = 50 µm. Arrowheads indicate necrotic/fibrotic muscle. **B.** Percent of centralized myonuclei out of 600 myofibers/TA comparing WT and *Dock3* KO mice. (n=5, student’s t-test, two-tailed, ****p<0.0001). **C.** Comparison of WT and *Dock3* KO mice fiber size distribution curve based on cross-sectional area (µm^2^). (n=5; two-way ANOVA with Tukey’s correction; ***p<0.001, ****p<0.0001). **D.** Percent of centralized myonuclei out of 600 myofibers/TA comparing *Dock3* alleles on *mdx*^*5cv*^ genetic background. (n=5, one-way ANOVA with Tukey’s correction; **p<0.01, ****p<0.0001). **E.** Comparison of fiber size distribution curve based on cross-sectional area (µm^2^) among *Dock3* alleles on *mdx*^*5cv*^ genetic background. (n=5; two-way ANOVA with Tukey’s correction; *p<0.05, ****p<0.0001).

### Adult Dock3 knockout mice have lower body weight due to loss of lean muscle mass

Given our data demonstrating that loss of DOCK3 expression in normal and dystrophin-deficient muscle results in differential muscle effects, we further interrogated the role of DOCK3 expression in overall muscle health and metabolism. Adult *Dock3* KO mice had significantly lower body weight compared to WT mice at six months of age (**Figure 5A**). Quantitative Magnetic Resonance Imaging (QMRI) in *Dock3* KO mice revealed reduced lean mass (**Figure 5B**), while there was no effect on overall fat mass (**Figure 5C)**. Lean muscle mass measurements were significantly lower in *Dock3* KO mice compared to WT controls (**Figure 5B**). As muscle is the primary source of glucose and given our previous findings linking DOCK3 to the PTEN/AKT muscle growth pathway, we next measured the ability of *Dock3* KO mice to respond to a glucose challenge via a glucose tolerance test (GTT)^32^. The glucose tolerance test revealed that *Dock3* KO mice had a significantly impaired response to the glucose bolus, indicating a deficit in glucose metabolic processing (**Figure 5D**). We then investigated the role of *Dock3* in global locomotor function using open field activity tracking to record basal activity levels in both aged-matched (6-month-old) WT and *Dock3* KO male mice. *Dock3* KO mice demonstrated significantly decreased distance traveled and average velocity compared to WT mice, indicating an overall reduction in basal locomotor function (**Figures 5E, 5F, Supplemental Figure 2**). This demonstrates an overall decrease in locomotor ability in *Dock3* KO mice at 6 months of age similar to that reported for the *mdx* mouse model and consistent with our findings that *Dock3* KO mice have significant muscle weakness and myofiber atrophy^33,34^,^35-40^. Additionally, we assessed cardiac function in WT and *Dock3* KO mice and found no significant differences in functional parameters (**Supplemental Figure 3**). These findings implicate a crucial role for DOCK3 in the maintenance of muscle mass and overall metabolic function in skeletal muscle.

**Figure 5.**
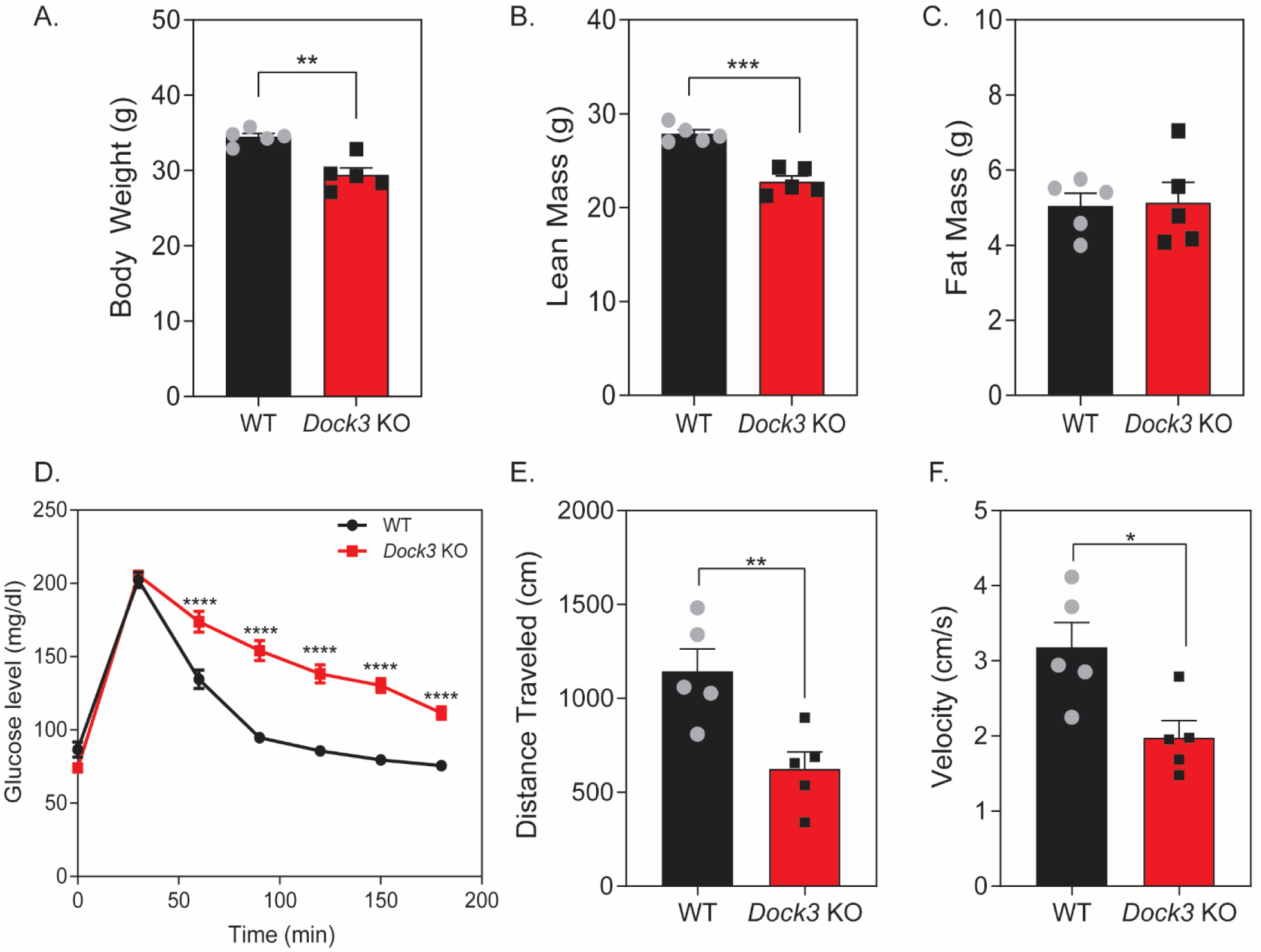
*Dock3* KO mice have abnormal muscle mass, metabolism, and overall decreased basal activity. **A.** Compared to WT mice, *Dock3* KO mice were significantly lower in gross body weight, **B.** had significantly decreased lean mass and, **C.** had no difference in fat mass observed. (n=5; student’s t-test, two-tailed; **p<0.01, ***p<0.001). **D.** Glucose tolerance test (GTT) revealed *Dock3* KO mice are glucose intolerant. (n=5; two-way ANOVA with Tukey’s correction; ****p<0.0001). **E.** *Dock3* KO mice had significantly decreased distance traveled (n=5; student’s t-test, two-tailed; **p<0.01). **F.** *Dock3* KO mice had significantly decreased average velocity (n=5; student’s t-test, two-tailed; *p<0.05).

### RNA-sequencing of Dock3 KO muscles reveals dysregulated muscle transcripts

To determine what pathways may be affected by the loss of DOCK3 protein in muscle, we performed RNA-sequencing (RNA-seq) on adult TA skeletal muscles from WT and *Dock3* KO mice. Transcriptome analysis revealed 280 differentially expressed RNA transcripts as shown by volcano plot (157 downregulated in blue and 123 upregulated in orange) (**Figure 6A**). Gene Ontology (GO) analyses revealed significant alterations in muscle structural and metabolic signaling pathways in the *Dock3* KO mice (**Figure 6B**), which is consistent with the severe *Dock3* KO phenotype observed. We identified the top 10 up- and down-regulated transcripts, excluding undefined transcripts, of *Dock3* KO mice compared to WT by sorting top genes according to the statistical significance (adjusted p-value) and strength of fold change induced (log_2_ fold change) (**Figures 6C, 6D**). We confirmed a significant decrease in *Dock3* mRNA transcript in the *Dock3* KO mouse muscles (**Figure 6D**). These findings reveal significant transcriptomic changes in important muscle structural and metabolic signaling pathways in the absence of DOCK3 protein.

**Figure 6.**
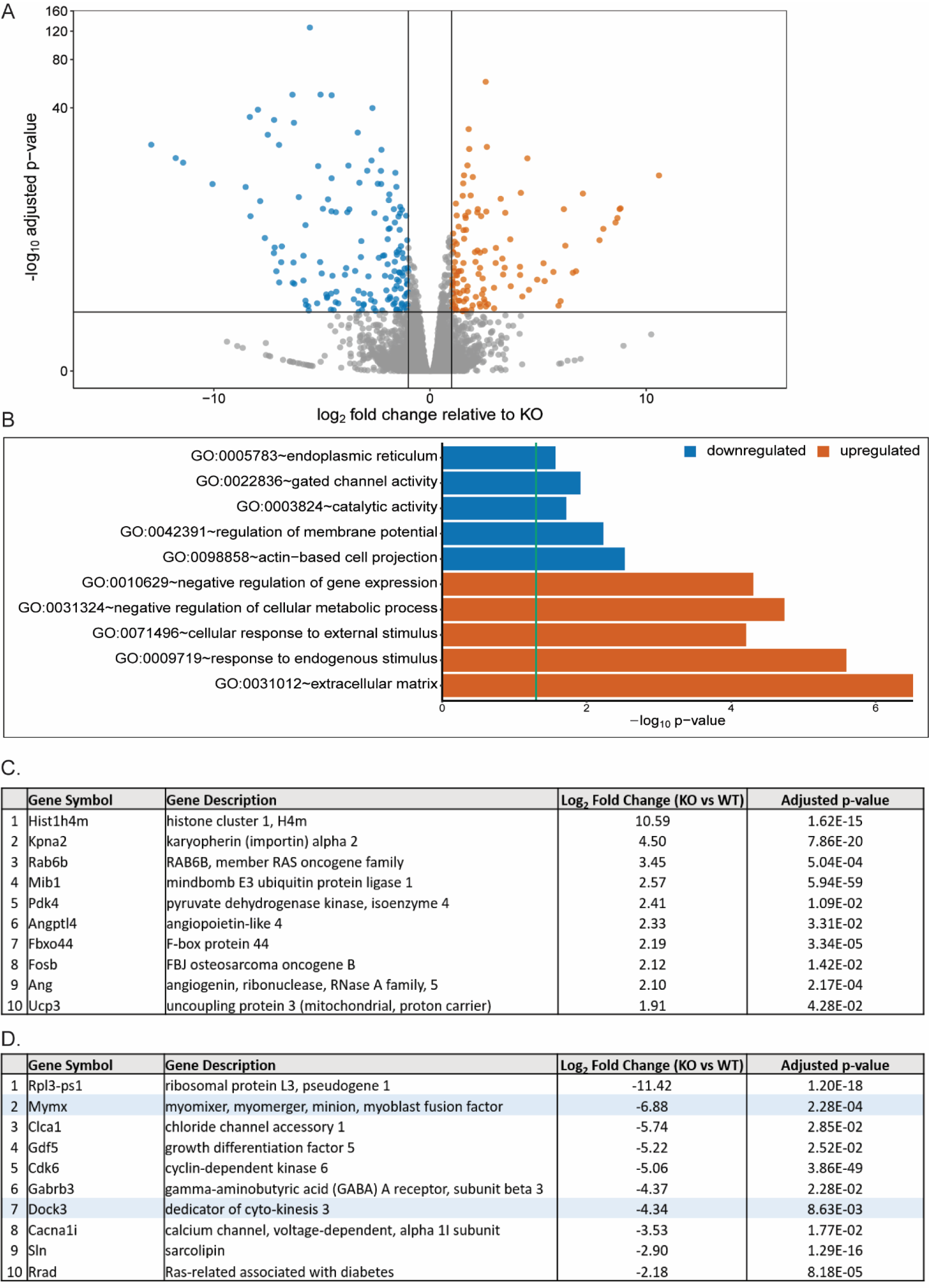
RNA-sequencing analysis of *Dock3* KO TA muscles reveals numerous dysregulated mRNA transcripts and signaling pathways. **A.** Summary graph of differentially regulated genes, with blue representing downregulated genes and orange representing upregulated genes relative to KO. **B.** Top downregulated and upregulated gene ontology terms as determined by Gene Ontology (GO) analyses compared to WT. **C.** Top ten upregulated genes in *Dock3* KO mice compared to WT. **D.** Top ten downregulated genes in *Dock3* KO mice compared to WT. Highlighted in blue are specific genes of interest.

### Dock3 KO myoblasts fail to differentiate and have reduced myogenic fusion capabilities

Previous studies have demonstrated that DOCK1 and DOCK5 are essential for normal myoblast fusion and subsequently differentiation via an actin nucleation factors^7,8,41^. Additionally, previous studies have demonstrated that DOCK3 interacts with the WAVE complex to activate RAC1, modulate actin dynamics, and promote cellular outgrowth^42^. We postulated that DOCK3’s function in modulating actin dynamics may influence muscle differentiation and fusion capabilities. We tested whether lentiviral shRNA knockdown of *DOCK3* mRNA might affected myogenic differentiation capabilities in primary human muscle cells. Healthy human myoblasts were treated with either mock, shRNA scrambled (negative control) or shRNA *DOCK3* and assessed for myogenic differentiation capability using myosin heavy chain (MyHC) fluorescence (**Figure 7A**). Myoblasts treated with shRNA *DOCK3* had significantly impaired myogenic differentiation compared to other control groups as evidenced by decreased amounts of MyHC-positive cells (**Figure 7B**). Western blot analysis validated the reduction of DOCK3 protein levels by shRNA *DOCK3* mRNA knockdown (**Figure 7C**). Moreover, we replicated this experiment using primary myoblasts isolated from our *Dock3* KO mice and WT mice and measured their myogenic differentiation capacity. We isolated primary myoblasts from adult WT and *Dock3* KO mouse TA muscles and evaluated their ability to fusion upon exposure to reduced serum medium. *Dock3* KO myoblasts showed poor myogenic differentiation capacity with an overall reduced amount of myotubes compared to aged-matched WT controls (**Figures 7D, 7E**). Taken together these studies demonstrate a critical role of DOCK3 protein in proper myogenic differentiation and myotube formation.

**Figure 7.**
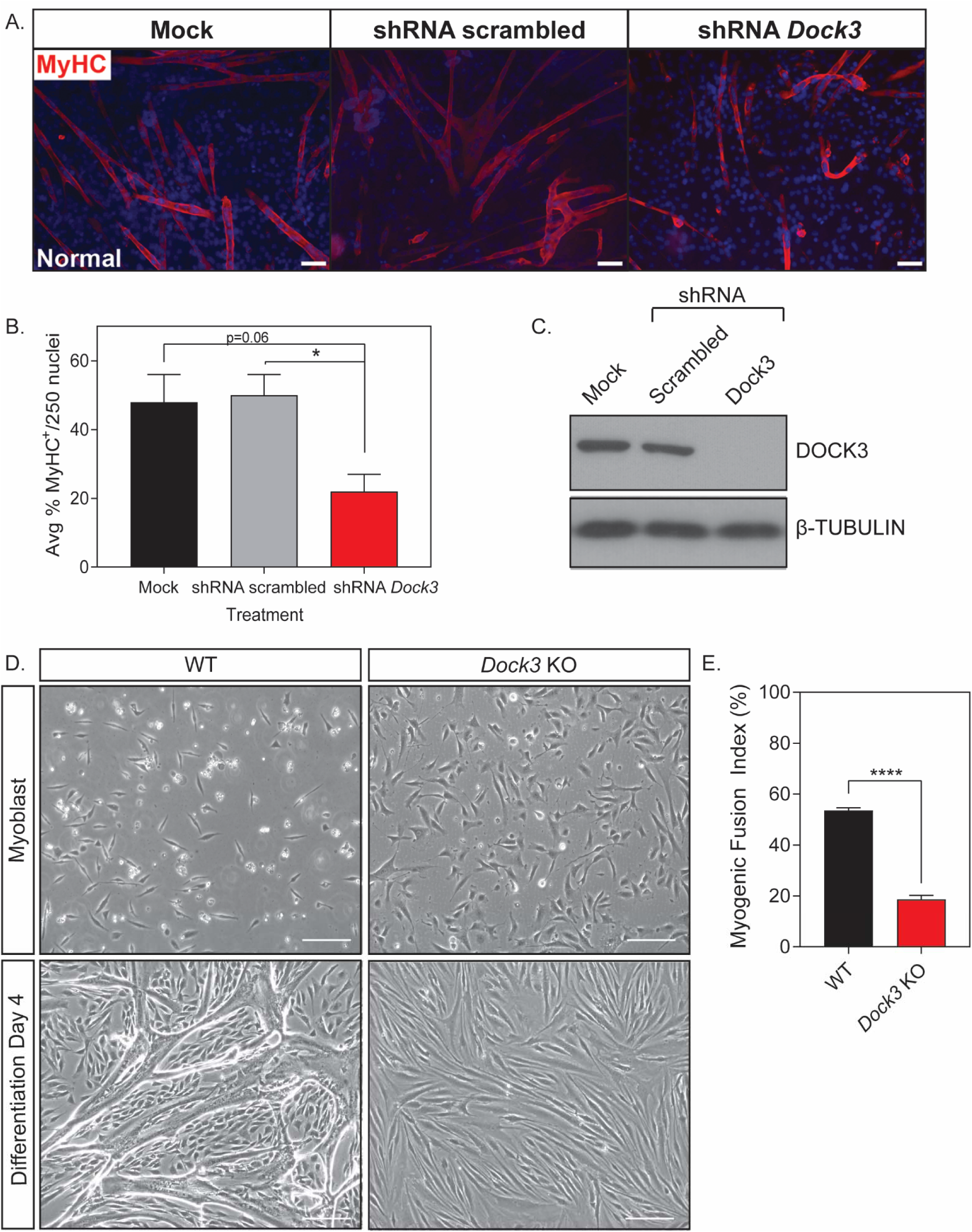
DOCK3 is essential for normal myoblast fusion. **A.** Myosin heavy chain staining reveals that knockdown of DOCK3 protein in normal human primary myotubes decreases myogenic fusion indices. MyHC (MF-20 antibody, DSHB Iowa, red) immunofluorescence of day 7 differentiated primary myotubes along with DAPI (blue). MOCK, shRNA scrambled, and shRNA *DOCK3* lentiviral particles were transduced at a MOI (multiplicity of infection) of 10. Scale bar = 50 µm. **B.** Average myogenic fusion indices as calculated by percentage of nuclei inside of MyHC^+^ cells out of 250 nuclei as previously described^63^. Myotubes with shRNA *DOCK3* knockdown have significantly less myogenic fusion than shRNA scrambled control (n=3 experimental replicates; one-way ANOVA with Tukey’s correction; *p<0.05). **C.** Western blot analysis of whole cell lysates taken from normal human day 7 differentiated myotubes transduced with mock (control), scrambled (shRNA internal control) or shRNA *DOCK3* knockdown inhibitor lentiviral particles showing successful knockdown of DOCK3 protein expression. **D.** Representative images of WT and *Dock3* KO mouse myoblasts undergoing differentiation. **E.** Quantification of myogenic fusion index where *Dock3* KO mouse myoblasts has significantly less myogenic fusion than WT mouse myoblasts. Scale bar = 100 µm. (n=3; student’s t-test, two-tailed; ****p<0.0001).

### Overexpression of myomixer (Mymx) modestly increases myogenic differentiation in Dock3 KO myoblasts

As we previously noted a significant decrease in expression levels of the myogenic fusion peptide myomixer (*Mymx*-*Myomerger*-*Minion*) in *Dock3* KO TA muscles (**Figure 7D**), we postulated that this decrease in myomixer expression may be the cause of the limited myogenic differentiation capacity of the *Dock3* KO myoblasts. We did not detect any binding interaction between DOCK3 and myomixer via co-immunoprecipitation studies (data not shown), thus we postulated that the decrease in myomixer expression may indirectly affect the ability of DOCK3 in an independent mechanism. We overexpressed FLAG-tagged *Mymx* (*Mymx* OE) or GFP in *Dock3* KO myoblasts, as validated via western blotting for the FLAG epitope in our *Mymx* plasmid (**Figure 8A, 8B**). The *Dock3* KO myoblasts transfected with *Mymx* OE demonstrated a slightly increased percentage of multi-nucleated cells as observed by increase in the myogenic fusion index compared to the GFP-control. No change in the percentage of myosin heavy chain-positive (MyHC^+^) nuclei or total nuclei was observed in the differentiated *Dock3* KO myoblasts (**Figure 8C-E**). We then assessed hallmark myogenic differentiation factors in the transfected cells via quantitative PCR (qPCR). The *Mymx* OE *Dock3* KO myoblasts had decreased expression of the differentiation factor *MyoG* and increased in *MyoD* expression (**Figure 8F, 8G**). No difference in the expression levels of *Myf5, Pax3*, or *Pax7* expression between the *Mymx* OE and GFP control groups (**Figure 8H-J**). Taken together, these results indicate that overexpression of myomixer has minimal effects in improving the myogenic differentiation capability in the *Dock3* KO myoblasts in *Dock3* KO myoblasts.

**Figure 8.**
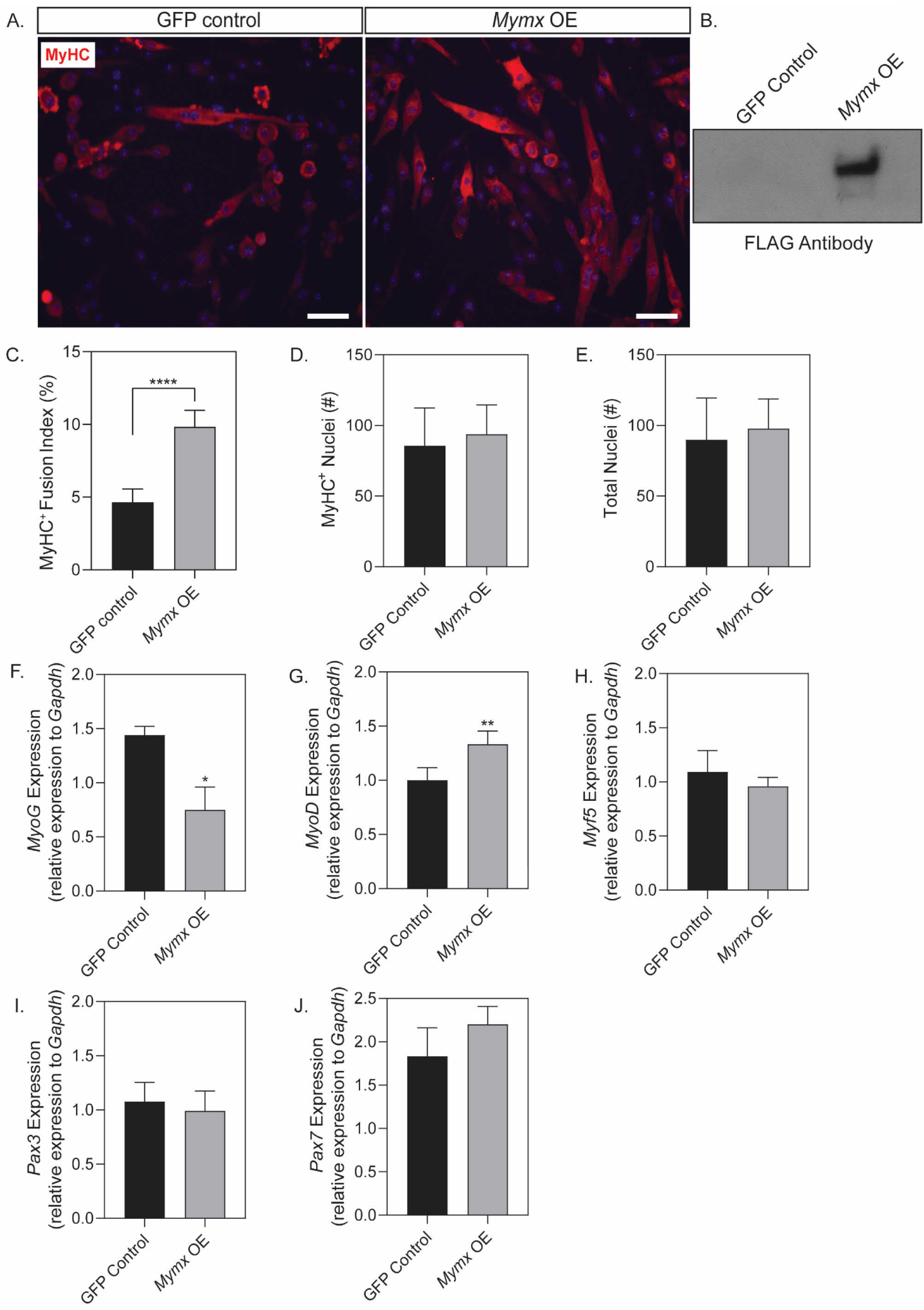
Myomixer overexpression in *Dock3* KO myoblasts induces myogenesis. **A.** Representative images of myoblasts transfected with either GFP control or myomixer (*Mymx*) overexpression (*Mymx* OE). **B**. Western blot for FLAG tag on *Mymx* plasmid to validate successful overexpression of *Mymx* via nucleofection. **C.** *Dock3* myoblasts transfected with *Mymx* OE have significantly higher number of MyHC^+^ multinucleated myotubes as seen by increased fusion index. Scale bars = 50 µm. (n=10, student’s t-test, two-tailed; ***p<0.001). **D.** No significant difference in number of MyHC^+^ nuclei (n=10, student’s t-test, two-tailed). **E.** No significant difference in total number of nuclei (n=10, student’s t-test, two-tailed). **F-J.** Quantitative-PCR (qPCR) of fusion and differentiation-related transcripts. Myoblasts with *Mymx* OE had significantly reduced expression of myogenin (*MyoG*) and increase in *MyoD*, but no change observed in *Myf5, Pax3*, or *Pax7* expression. (n=5, student’s t-test, two-tailed, *p<0.05, **p<0.01).

## Discussion

We have identified *DOCK3* as a modifier of normal and dystrophin-deficient skeletal muscle health and function. First, we showed that DOCK3 is strongly elevated in expression levels in dystrophin-deficient mouse skeletal muscles consistent with the DOCK3 expression profile in human patients^8^. While complete loss of *Dock3* expression in *Dock3* KO and *Dock3:mdx*^*5cv*^ DKO mice have detrimental effects on skeletal muscles, haploinsufficiency of *Dock3*^*+/-*^:*mdx*^*5cv*^ improves overall myofiber size and skeletal muscle pathology (**Figure 9**). These findings were further validated in the zebrafish *sapje* DMD zebrafish. Taken as a whole, these studies demonstrated that the reduction of *Dock3* expression in DMD models prevents the severity of dystrophic muscle pathology. These findings potentially represent the discovery of DOCK3 as new therapeutic target for DMD, as previous findings have identified factors that provide protective haploinsufficiency in *mdx* mice. For example, the haploinsufficiency of the transcription factors *Six4* and *Six5* in *mdx* mice similarly resulted in improved muscle grip strength and extended life span when haploinsufficient in *mdx* mice^43^.

**Figure 9.**
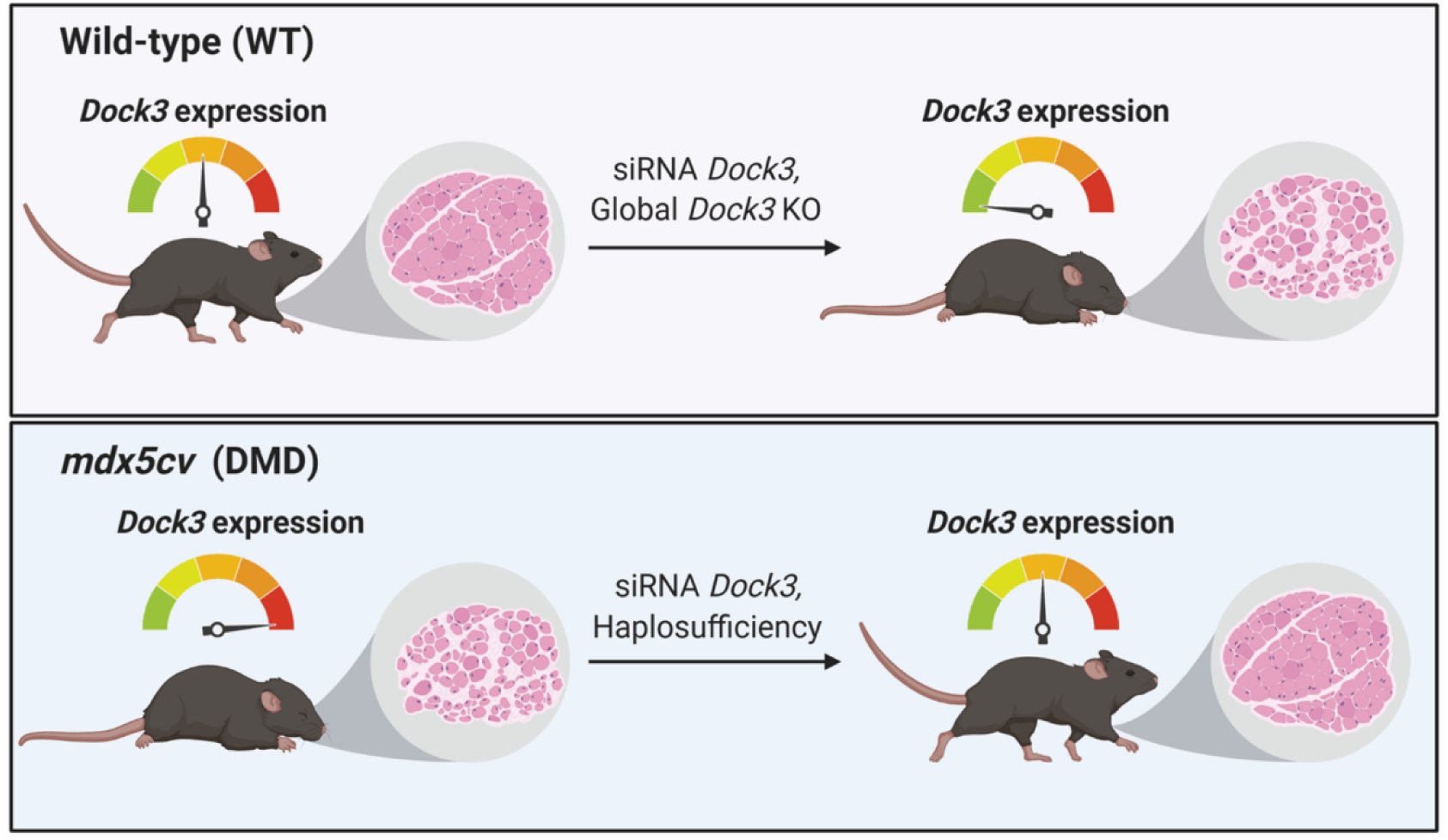
DOCK3 dosage is tightly regulated in normal and dystrophic muscles. In normal muscle, loss of DOCK3 results in worsened muscle phenotype and impairment of myogenic fusion. In dystrophic muscle that has significantly heightened expression of DOCK3, reduction of DOCK3 results in improved muscle health and function. However, total loss of DOCK3 in dystrophic muscle worsens muscle phenotype, demonstrating the critical gene-dosage effect of DOCK3.

Complete loss of DOCK3 expression in both wild type and dystrophin-deficient muscle resulted in overall worse muscle health, decreased myoblast fusion, glucose intolerance, and overall defective muscle locomotive function. This is consistent with the reporting of clinical muscle hypotonia and overall muscle weakness in patients with pathogenic *DOCK3* variants^10-12^. While the roles of other DOCK proteins in muscle are not well described, it is possible that DOCK3 may have some overlapping and non-overlapping functions with DOCK4 based on similar protein sequence structure and domains.

Additionally, we observed multiple myogenic differentiation factors that were decreased in expression levels in the *Dock3* KO muscles via transcriptomic pathway analysis. A key question remains on whether DOCK3 functions to alter the filamentous actin in myoblasts or works through another mechanism to regulate the myogenic differentiation process. Our studies revealed that overexpression of myomixer in *Dock3* KO myoblasts while slightly increasing the myogenic differentiation capability, did not fully restore proper myogenic differentiation. It remains likely that DOCK3 has essential interactions with actin and other key molecules that may be the source of the impaired myogenic differentiation observed in the *Dock3* KO myoblasts.

The mechanism of action of DOCK3 in myogenesis and its role as a guanine nucleotide exchange factor (GEF) remains of key interest to understand its function in muscle disease. As a GEF, DOCK3 activates RAC1, a Rho-GTPase, which has been shown to be tightly regulated in myogenesis^44^. We previously demonstrated that DOCK3 overexpression failed to activate RAC1 in DMD myotubes^14^. Other studies have demonstrated that RAC1 cannot activate in atrophied myofiber membranes when the dystrophin-associated complex (DGC) is structurally compromised^45^. Overexpression of a constitutively active RAC1 in C_2_C_12_ mouse skeletal myoblasts inhibits proper differentiation demonstrating the importance of RAC1 activation status in regulating temporal signals in myogenesis^46^. Similar to the glucose impairment in the *Dock3* KO mice, the muscle *Rac1* KO mice show impaired glucose processing^47^. Interestingly it has been shown that overexpression of Def-6 (also called HMGB1), a GEF protein similar to DOCK3, impairs the downregulation of RAC1 activity and results in inhibited myogenic differentiation^48^. These studies provide mechanistic insight into the molecular interplay between DOCK3 and RAC1 in dystrophin-deficient muscles and myogenesis.

Our findings suggest that DOCK3 plays a key role in normal muscle fusion and affect dystrophin-deficient muscle pathologies and outcomes. Furthermore, as DOCK3 expression levels are tightly correlated with dystrophic disease progression, DOCK3 may be a potential DMD biomarker that could be used to measure disease severity in muscle. Moreover, our findings also provide strong evidence for DOCK3 as a potential novel therapeutic target for the treatment of DMD. Future experiments aimed at further elucidating the mechanism of DOCK3 in muscle differentiation, signaling, and its contribution towards muscle metabolic regulation are warranted.

## Materials and Methods

### Mice

*Dock3* global knockout mice (Jax mice stock# 033736) were obtained from the laboratory of Dr. David Shubert (Salk Institute) and have been previously described^9^. These mice contain a *beta-galactosidase* cassette integrated in-frame into exon 2 of mouse *Dock3* locus resulting in a complete disruption of the gene. Wild type (*C57BL/6J*; stock# 000664) and *mdx*^*5cv*^ (stock# 002379) were originally obtained from Jackson Labs (Bar Harbor, ME). All mice were maintained on the *C57BL/6J* strain background. All mouse strains were maintained under standard housing and feeding conditions with the University of Alabama at Birmingham Animal Resources Facility under pathogen-free, sterile conditions under the animal protocol number 20232.

### Zebrafish morpholino experiments

Wild type *AB* and *sapje* (*dmd*^*ta222a*^ mutation on the *AB* strain background) zebrafish were used for all zebrafish morpholino experiments. A *dock3* morpholino (5’-GCCTCAGATCAATCAACTCGTTCAT-3’) and a control FITC-labeled (5’-CCTCTTACCTCAGTTACAATTTATA-3’) non-targeting morpholino (Gene Tools LLC; Philomath, OR) were used for all morpholino injections. Zebrafish morpholinos were injected into one-cell embryos obtained from *sapje* heterozygote matings at amounts of 0.1, 0.5, 1, 2, 3, and 6 nanograms (ng) in a solution of 1x Danieau buffer, water, and phenol red (Sigma Aldrich, St. Louis, MO) as previously described^49^. For myosin heavy chain (MyHC) zebrafish larvae were fixed in 4% paraformaldehyde (Electron Microscopy Sciences) overnight at 4°C and then following 3x washes in 1xPBS for five minutes, were incubated for 1 hour at room temperature with F-59 (MyHC; Developmental Studies Hybridoma Bank, Iowa City, IA) in the dark. Larvae were then washed 3x in 1xPBS for five minutes per wash, and then incubated with anti-mouse Alexa Fluor-568 (Invitrogen; Cat# A-11004) for 45 minutes at room temperature in the dark. After an additional three washes in 1x PBS, the fish were then placed on superfrosted slides (FisherScientific; Hampton, NH) and imaged on a Nikon Eclipse E-1000 microscope. All adult fish were fed a standard diet of Artemia salina (brine shrimp) three times per day under a 14 hour on, 10 hour off light cycle in 3L tanks with a density of no more than 20 fish per tank as standard of care guidelines. All zebrafish strains were maintained University of Alabama at Birmingham Animal Resources Aquatics Facility under pathogen-free conditions under the animal protocol number 20230.

### Zebrafish muscle birefringence assay

Subsequently following the locomotor assay, birefringence assay was performed on 5 dpf larvae as previously described. Larvae were anesthetized using MS-222 and placed on microscope with polarizing light attachment. In this assay, when the polarized light is shone on the larvae, a white light is refracted back and captured in the camera if the muscle is healthy and has organized fibers. However, if the muscle is broken down and disorganized (worsened muscle phenotype), the polarized light will shine straight through and will appear in the camera as black (absence of white refracted light).

### In situ hybridization (ISH) zebrafish experiments

The zebrafish *dock3* mRNA (*dock3-201* ENSEMBLE transcript; ENSDARG00000063180) was used as a template and an antisense region containing 300 base pairs sequence of the end of *dock3* mRNA coding sequence was used for *in vitro* transcription probe synthesis. *In vitro* transcription reactions performed using the MegaScript Sp6 (Cat# AM 1330) and MegaScript T7 Transcription (Cat# AM 1334) kits (ThermoFisher Scientific) were used following the manufacturer’s guidelines. Morpholino injected zebrafish embryos were placed in fish water containing 1x PTU (Sigma; Cat# P7629) at twenty-four hours post fertilization. *In situ* hybridization reactions were performed on 4% paraformaldehyde (Electron Microscopy Sciences; Hatfield, PA; Cat# 15710) zebrafish embryos using the DIG RNA Labeling Kit (MilliporeSigma; Burlington, MA; Cat# 11175025910) following a standardized protocol^50^. The DIG-labeled embryos were imaged under the dissection scope (Nikon, SMZ1500; Tokyo, Japan) with EXFO Fluorescence illumination system, X-Cite 120 (Photonic Solutions Inc.; Edinburgh, UK), using a Nikon Eclipse E-1000 microscope attached to a Hamamatsu digital camera, and images were acquired using Openlab software version 3.1.5 (Improvision; PerkinElmer; Waltham, MA).

### Human muscle samples

Primary *vastus lateralis* muscle biopsies from normal and DMD patients were obtained from consented patients under the approved Boston Children’s Hospital protocol 03-12-205 and described elsewhere^14,15^. Additional samples were collected under the UAB protocol 300002164. All patient samples were de-identified and secondary confirmation of DMD mutations were performed using MLPA analysis and/or exome sequencing.^51^

### Mouse Activity Tracking

Mouse overall locomotive activity measurements were performed as previously described^52^. Twenty-four hours prior to experiment termination and tissue harvest, mice were analyzed for overall locomotive activity using the Ethovision XT software platform (Noldus; Leesburg, VA) with isolated individual chambers adapted from a previously described protocol^14^. Mice were adapted to the room and open-field chambers one day prior to activity and were given a 5-minute additional adaptation period prior to activity recording. Mouse activity was recorded for six minutes with no external stimulation.

### Myofiber and myogenic fusion calculations

The cross-sectional area (CSA) of the skeletal muscle sections was calculated in a manner previously described^14^. Briefly, 600 TA myofibers were counted and cross-sectional area (µm^2^) was measured via a series of overlapping H&E microscopy images, and quantified in Fiji software platform^53^. Myogenic fusion indices were determined by immunofluorescent staining using the MF-20 (myosin heavy chain/MyHC) as a marker of myogenic differentiation. Fusion of myoblasts was determined by the detection of more than one nuclei within an MyHC+ myotube.

### DEXA Quantitative Magnetic Resonance (QMR) Imaging

To evaluate body composition (fat and lean tissue mass) *in vivo*, 6-month male WT and *Dock3* KO mice (5 mice/genotype) were measured using the EchoMRI™ 3-in-1 composition analyzer (software version 2016, Echo Medical, Houston, TX). Individual fat and lean mass measurements were measured in grams (g) and were analyzed using student’s t-test between WT and *Dock3* KO mice.

### Glucose tolerance test (GTT)

Mice were fasted for 16 hours prior to afternoon administration of a bolus of D-glucose (MilliporeSigma; Cat# G8270) was intraperitoneal (IP) injected at a concentration of 3 mg/gram of mouse body weight. Blood glucose was measured on a commercially obtained glucometer (Nipro Diagnostics; Ft. Lauderdale, FL) using 10 µl of whole serum from tail bleeds placed on standardized glucose meter test strips.

### siRNA knockdown in cardiotoxin-injured muscles

Adult male *mdx*^*5cv*^ mice (4-6 months old) were injected in their TA skeletal muscles with cardiotoxin (10 µM) and siRNA pooled oligos (10 µg; GE Healthcare Dharmacon Inc; Lafayette, CO) containing either scrambled siRNAs or siRNAs targeting mouse *Dock3* mRNA following a protocol previously established^54^. The contralateral TA muscle was used as a sham phosphate-buffered saline (ThermoFisher Scientific; Waltham, MA; Cat#10010049) control. Seven days following injections, mice were euthanized, and their TA skeletal muscles were slow-frozen in TissuePlus O.C.T (FisherScientific; Hampton, NH; Cat#23-730-571), for histological and molecular analysis.

### Immunofluorescence and immunohistochemistry

Mouse skeletal muscles were cryo-frozen in a TissuePlus O.C.T (FisherScientific; Hampton, NH; Cat#23-730-571) Isopentane (FisherScientific; Cat# AC397221000) liquid nitrogen bath as unfixed tissues. Mouse hearts were perfusion fixed in 10% neutral buffered formalin (MilliporeSigma) overnight at 4°C was performed before embedding the tissue sample into paraffin blocks. Blocks were later cut on a cryostat and 7-10 µm thick sections were placed on SuperFrost Plus Gold slides (ThermoFisher; Cat# FT4981GLPLUS). Hematoxylin and eosin (H&E) staining was performed as previously described^15^. For immunofluorescent staining, following de-paraffinization the slides were incubated in eBioscience IHC Antigen Retrieval Solution (ThermoFisher; Cat# 00-4955-58) and washed in 1x PBS three times for five minutes, and then incubated with blocking reagent from the Mouse-on-Mouse (M.O.M) kit (Vector Laboratories; Burlingame, CA; Cat# BMK-2202). Slides were incubated for 1 hour at room temperature in primary antibody.

### Western Blot

Protein lysates were obtained from either pestle-homogenized tissues or cell lysates in M-PER lysis buffer (ThermoFisher; Cat# 78501) supplemented with 1x Complete Mini EDTA-Free protease inhibitor cocktail tablets (Roche Applied Sciences; Cat# 04 693 159 001; Penzburg, Germany). Protein lysates were quantified using a Pierce BCA Protein Assay Kit (ThermoFisher; Cat# 23225). Unless stated otherwise, fifty micrograms of whole protein lysate was used for all immunoblots, and electrophoretically resolved on 4-20% Novex tris-glycine gels (ThermoFisher: Cat# XPO4205BOX). Protein samples were then transferred to 0.2 µm PVDF membranes (ThermoFisher: Cat# LC2002), blocked in 0.1x TBS-Tween in 5% non-fat milk for 1 hour before being incubated in primary antibody overnight at 4°C with gentle rocking. Blots were washed in 0.1% TBS-Tween 3 times for 5-minute intervals before being incubated with secondary antisera (mouse or rabbit) conjugated to HRP for 1 hour at room temperature with gentle agitation. Membranes were then washed in 0.1% TBS-Tween 3 times for 5-minute intervals before being incubated with secondary antibodies (either mouse or rabbit) conjugated to horse radish peroxidase (HRP) for 1 hour at room temperature with gentle agitation. Following another 3 washes for 15-minute intervals at room temperature, membranes were then treated with RapidStep ECL Reagent (MilliporeSigma; Cat# 345818-100ML) and exposed to x-ray film (Genesee Scientific). Some Western blot images were acquired using a Typhoon Variable Mode Imager (Amersham Pharmacia; Little Chalfont, UK). Some membranes were stripped using Restore Plus Western Blot Stripping Buffer (ThermoScientific; Cat# 46428) and later probed with a different primary antibody.

### RNA-sequencing and data analyses

Adult male 6-month old wild type and *Dock3* KO mouse TA muscles (n = 3 muscles per genotype cohort) were biopsied and total RNA was extracted following mechanical homogenization using the GenElute Total RNA Purification Kit (MilliporeSigma; Cat# RNB100-50RXN) following the manufacturer’s guidelines. The total RNA was amplified using the Sure Select Stranded RNA-Seq kit (Agilent Technologies; Santa Clara, CA) using standard protocols. A Ribominus kit (ThermoFisher; Cat# K155002) was used to deplete large ribosomal RNAs. The sequencing was done on the Illumina HiSeq2500 with paired end 50bp reads following the manufacturer’s protocols by the UAB Genomics Core Facility.

All samples contained a minimum of 17.7 million reads with an average number of 19.8 million reads across all biological replicates. The FASTQ files were uploaded to the UAB High Performance Computer cluster for bioinformatics analysis with the following custom pipeline built in the Snakemake workflow system (v5.2.2)^55^: first, quality and control of the reads were assessed using FastQC, and trimming of the bases with quality scores of less than 20 was performed with Trim_Galore! (v0.4.5). Following trimming, the transcripts were quasi-mapped and quantified with Salmon^56^ (v0.12.0, with ‘--gencode’ flag for index generation and ‘-l ISR’, ‘--gcBias’ and ‘-- validateMappings’ flags for quasi-mapping) to the mm10 mouse transcriptome from Gencode release 21. The average quasi-mapping rate was 70.4% and the logs of reports were summarized and visualized using MultiQC^57^ (v1.6). The quantification results were imported into a local RStudio session (R version 3.5.3) and the package “tximport”^58^ (v1.10.0) was utilized for gene-level summarization. Differential expression analysis was conducted with DESeq2^59^ package (v1.22.1). Following count normalization, principal component analysis (PCA) was performed and genes were defined as differentially expressed genes (DEGs) if they passed a statistical cutoff containing an adjusted p-value <0.05 (Benjamini-Hochberg False Discovery Rate (FDR) method) and if they contained an absolute log_2_ fold change >=1. Functional annotation enrichment analysis was performed in the NIH Database for Annotation, Visualization and Integrated Discovery (DAVID, v6.8) by separately submitting upregulated and downregulated DEGs. A p-value <0.05 cutoff was applied to identify gene ontology (GO) terms. The FASTQ files of the current study have been uploaded to NCBI’s Gene Expression Omnibus under accession number GSE141621.

### Real time quantitative PCR (rt-qPCR)

Total RNA was extracted using the miRVana (ThermoFisher; Cat# AM1560) kit following the manufacturer’s protocol. Total RNA was reverse transcribed using the Taqman Reverse Transcription kit following the manufacturer’s protocol (Applied Biosystems; Foster City, CA; Cat# N8080234). Taqman assay probes were all purchased from Applied Biosystems corresponding to each individual gene. Quantitative PCR (qPCR) Taqman reactions were performed using Taqman Universal PCR Master Mix (Applied Biosystems; Cat# 4304437). Samples were run on the Fluidigm Biomark HD platform (Fluidigm Corp.; San Francisco, CA) in 96.96 Dynamic Array plates. Relative expression values were calculated using the manufacturer’s software and further confirmed using the 2^−ΔΔCt^ method^60^.

### Muscle Myoblast Cell Cultures

Primary muscle cells were isolated from postnatal seven-day-old WT or *Dock3* KO mouse hindlimbs via a pre-plate purification strategy for enrichment of a slow-adhering purification as previously described^61,62^. Primary muscle cells were later evaluated for myogenic capacity via immunofluorescent staining of desmin to ensure for a myogenic population greater than 95%. Muscle cells were grown in Skeletal Muscle Growth Medium (Promocell; Heidelberg, Germany; Cat# C-23060) supplemented with 1x antibiotic-antimycotic (ThermoFisher Scientific; Cat# 15240062) and 1 ng/10ml rhFGF (Promega; Madison, WI; Cat# G5071). Cells were seeded at a density of 1.5×10^6^ per 10 cm^2^ plate or 2-chamber slides (Corning Inc.; Corning, NY) coated with 1% rat tail collagen (Millipore Sigma; Cat# 08-115). When cells reached 90% confluency, they were switched to differentiation medium, which is composed of DMEM GlutaMAX™, 1% of Antibiotic-Anti-mycotic (Invitrogen: Cat# 1520), and Promocell Human Skeletal Muscle Differentiation Media (Cat# C-23061). Myogenic fusion assays and immunofluorescent labeling was performed as previously described^63^.

### Nucleofection of Myomixer fusion protein in mouse Dock3 KO Primary Myoblasts

Myomixer-FLAG was packaged in the pLVX-TetON (Takara Bio; Mountain View, CA) plasmid and have been described in previous literature^64^. The plasmid was subcloned into pIRES-1a-hrGFP (Stratagene; San Diego, CA) plasmid. Nucleofection of *Dock3* KO primary myoblasts in culture was done according to the optimized protocol for skeletal muscle myoblasts following the manufacturer’s protocol (Lonza; Basel, Switzerland). Briefly, myoblasts were grown in their respective culture media in 100 cm^2^ plates until they reached 80% confluency, corresponding to approximately 1.5 × 10^6 cells. The medium was removed from the plate and cells were trypsinized with 1 ml of trypsin-EDTA (Invitrogen) to detach them completely. Subsequently, 4ml of medium was added to the plate and then the cells and the culture medium were centrifuged at 1000 x g for 5 min at room temperature. The supernatant was then removed, and cell pellets were resuspended in 100 µL of Nucleofector solution (Lonza) in which either 2ug of myomixer plasmid or 2 ug pmaxGFP plasmid were added to respective cohorts. The cells were analyzed 24-48 hours after the nucleofection by fluorescence microscopy with an inverted Leica microscope. After 48 hours post-nucleofection, proliferation media was removed and washed with 1X PBS (Dulbecco) which was aspirated and replaced with differentiation media.

### Myogenic Fusion Assays

Myoblasts from *Dock3* global KO mice were nucleofected with myomixer-FLAG and its GFP control in four-well chamber slides at 100,000 cells/per well, separately (as described above). To assess myogenic fusion, cells were fixed at day 4 of differentiation with 4% paraformaldehyde for 20 minutes at 4°C followed by permeabilization with 0.1% triton™X-100 (Sigma Aldrich) for 30 minutes at room temperature and blocking in 1xPBS with 1% fetal bovine serum (Gibco). Cells were incubated with primary antibody for myosin heavy chain (MF-20—1:100, Developmental Studies Hybridoma Bank) overnight at 4°C, followed by three washes with 1x PBS and incubation with secondary antibody Alexa Fluor 555 (Invitrogen). Slides were washed and mounted with Vectashield® Mounting Media (Vector) with DAPI. Images were obtained in Zeiss 710 confocal system at 20X. The fusion index was calculated as follows: (MF20-stained myocytes containing ≥2 nuclei/total number of nuclei) × 100, as previously described^65^.

### Clustal Omega Alignment

CLUSTAL Omega alignments using freely available software (https://www.ebi.ac.uk/Tools/msa/clustalo/) as described^66^. Accession numbers for protein sequences: zebrafish (*Danio rerio*, XP_009294754.1), mouse (*Mus musculus*, NP_700462.2), and human (*Homo sapiens*, NP_004938.1).

### Cardiac functional measurements: *transthoracic echocardiography*

The Vevo 3100 (VisualSonics Inc, Canada) imaging system was used for echocardiography to evaluate individual parameters of cardiac structure and function *in vivo.* 6-month male WT and *Dock3* KO mice (5 mice/genotype) were anesthetized using 1-2% isofluorane continuously suppled from surgivet vaporizer and both long- and short-axis high resolution images were acquired for heart function measurements and analysis. Each individual parametric measurement was compared between WT and *Dock3* KO mice using a student’s T-test. (**Supplemental Figure 3**).

### Statistical Analyses

All graphs were represented as mean ± SEM. Unless otherwise mentioned, a two-tailed student’s t-test was performed for all single comparisons and either a one-way or two-way analysis of variance (ANOVA) with Tukey correction was performed for all multiple comparisons. GraphPad Prism version 8 software (GraphPad Software; San Diego, CA) was used for all statistical analyses. An *a priori* hypothesis of *p<0.05, **p<0.01, ***p<0.001, and ****p<0.0001 was used for all data analysis.

## List of Abbreviations

CSA: cross-sectional area
ctx: cardiotoxin
DAPC: dystrophin-associated complex
DEXA: dual energy x-ray absorptiometry
DIG: digoxigenin-UTP
DMD: Duchenne muscular dystrophy
DOCK: Dedicator of cytokinesis
dpf: days post fertilization
ECM: extracellular matrix
GEF: guanine nucleotide exchange factor
GFP: green fluorescent protein
GRMD: Golden retriever muscular dystrophy
GTT: glucose tolerance test
ISH: *in situ* hybridization
MLPA: multiplex ligation-dependent probe amplification
MOCA: modifier of cellular adhesion
MO: morpholino
MyHC: Myosin Heavy Chain
PTU: 1-Phenyl-2-Thiourea; N-Phenylthiourea
QMR: Quantitative Magnetic Resonance
qPCR: quantitative polymerase chain reaction
shRNA: short hairpin RNA
siRNA: small interfering RNA
TA: tibialis anterior

## Acknowledgements

We would like to thank Dr. David Schubert (Salk Institute) for the generous gift of the *Dock3* KO mice. We would like to thank Dr. Michael Crowley and the staff of the UAB Genomics Core Facility for RNA-sequencing processing and initial analyses. We would like to thank Eddie Bradley for the mouse cardiovascular function analyses. We would like to thank Drs. Timothy Nagy and Maria Johnson at the UAB Small Animal Phenotyping Core for DEXA measurements. We would like to thank Drs. David Schneider (UAB), Glenn Rowe (UAB), and Emanuela Gussoni (Boston Children’s Hospital) for critical reading of our manuscript. We would like to thank Dr. Susan Farmer and her staff for care and maintenance of our animals. Lastly, we would like to thank the patients and their families who contributed samples that were used in this study and helped to make this research possible.

## Manuscript Contributions

A.L.R., Y.W., R.M.H., A.S., and M.S.A. all performed experiments and analyzed data presented in this manuscript. L.I. and D.K.C. performed bioinformatics analyses of RNA-sequencing data. L.J.D. and G.V.H. analyzed and interpreted mouse cardiovascular data. M.A.L. and S.R.G. consented patients and obtained human biopsies. D.P.M. contributed novel reagents and analyzed data. T.V.G. analyzed and interpreted mouse locomotive activity data. A.L.R., Y.W., and M.S.A. analyzed all the data, and wrote the manuscript. All authors have fully approved of the final version of this manuscript.

## Supplemental Figure Legends

**Supplemental Figure S1. Clustal Omega Alignment of zebrafish, mouse, and human DOCK3 protein alignments.** Zebrafish (XP_009294754.1), mouse (NP_700462.2), and human (NP_004938.1) DOCK3 amino acid sequence alignments. Mice and zebrafish protein sequences demonstrate 98% and 85% sequence identity to human DOCK3 protein, respectively. Table showing homology DOCK3 amino acid conservation comparisons among all three species (excerpt from full-length protein analysis). CLUSTAL Omega alignments using freely available software (https://www.ebi.ac.uk/Tools/msa/clustalo/) as described^66^.

**Supplemental Figure S2. Representative tracings of basal activity in WT and *Dock3* KO mice.** Activity tracings of WT and *Dock3* KO adult 6-month-old male mice recorded for 5 minutes. No major differences were seen in trace patterns between cohorts.

**Supplemental Figure S3. *Dock3* KO mice have normal cardiac function.** *Dock3* KO mice were assessed using echocardiograms for various parameters of cardiac function, summarized in table. There were no significant differences in any functional parameter such as left ventricular ejection fraction (LV EF), LV fractional shortening (LV FS) or VCFr. There were some changes in parameters associated with volume of the heart, which correlates to the overall decrease in body size observed in *Dock3* KO mice (**Figure 5A**). When normalized to body size, there is no significant difference in any parameters. (n=5; student’s t-test, two-tailed; *p<0.05, **p<0.01, ***p<0.001). Abbreviations: IVSd, Interventricular septum thickness at end-diastole; IVSs, interventricular septum thickness at end-systole; LVEDD, left ventricular end-diastolic diameter; LVESD, left ventricular end-systolic diameter; LVPWd, left ventricular posterior wall thickness at end-diastole; LVPWs, left ventricular posterior wall thickness at end-systole, LV Vol;s, left ventricular volume at systole; LV Vol;d, left ventricular volume at diastole; LV EF, left ventricular ejection fraction; LV CO, left ventricular cardiac output; LV FS, left ventricular fractional shortening; VCFr, mean velocity of circumferential fiber shortening; HR, heart rate.

## References Cited

1. Ballarino, M., Morlando, M., Fatica, A. & Bozzoni, I. Non-coding RNAs in muscle differentiation and musculoskeletal disease. The Journal of Clinical Investigation 126, 2021–2030 (2016).

2. Cacchiarelli, D., et al. miR-31 modulates dystrophin expression: new implications for Duchenne muscular dystrophy therapy. EMBO Rep 12, 136–141 (2011).

3. Waldrop, M.A., Gumienny, F., Weiss, R.B. & Flanigan, K.M. Low-Level Dystrophin Expression Attenuating the Dystrophinopathy Phenotype. Muscle & Nerve, n/a-n/a (2017).

4. Peter, A.K. & Crosbie, R.H. Hypertrophic response of Duchenne and limb-girdle muscular dystrophies is associated with activation of Akt pathway. Experimental Cell Research 312, 2580–2591 (2006).

5. Dumont, N.A., et al. Dystrophin expression in muscle stem cells regulates their polarity and asymmetric division. Nat Med advance online publication(2015).

6. Laurin, M. & Côté, J.-F. Insights into the biological functions of Dock family guanine nucleotide exchange factors. Genes & Development 28, 533–547 (2014).

7. Moore, C.A., Parkin, C.A., Bidet, Y. & Ingham, P.W. A role for the Myoblast city homologues Dock1 and Dock5 and the adaptor proteins Crk and Crk-like in zebrafish myoblast fusion. Development 134, 3145–3153 (2007).

8. Laurin, M., et al. The atypical Rac activator Dock180 (Dock1) regulates myoblast fusion in vivo. Proceedings of the National Academy of Sciences 105, 15446–15451 (2008).

9. Chen, Q., et al. Loss of Modifier of Cell Adhesion Reveals a Pathway Leading to Axonal Degeneration. The Journal of Neuroscience 29, 118–130 (2009).

10. Helbig, K.L., Mroske, C., Moorthy, D., Sajan, S.A. & Velinov, M. Biallelic loss-of-function variants in DOCK3 cause muscle hypotonia, ataxia, and intellectual disability. Clinical Genetics, n/a-n/a (2017).

11. Iwata-Otsubo, A., et al. DOCK3-related neurodevelopmental syndrome: Biallelic intragenic deletion of DOCK3 in a boy with developmental delay and hypotonia. American Journal of Medical Genetics Part A 176, 241–245 (2018).

12. Wiltrout, K., et al. Variants in DOCK3 cause developmental delay and hypotonia. European Journal of Human Genetics 27, 1225–1234 (2019).

13. Wiltrout, K., et al. Variants in DOCK3 cause developmental delay and hypotonia. European Journal of Human Genetics (2019).

14. Alexander, M.S., et al. MicroRNA-486–dependent modulation of DOCK3/PTEN/AKT signaling pathways improves muscular dystrophy–associated symptoms. The Journal of Clinical Investigation 124, 2651–2667 (2014).

15. Alexander, M., et al. Regulation of DMD pathology by an ankyrin-encoded miRNA. Skeletal Muscle 1, 27 (2011).

16. Quinn, M.E., et al. Myomerger induces fusion of non-fusogenic cells and is required for skeletal muscle development. Nature Communications 8, 15665 (2017).

17. Bi, P., et al. Fusogenic micropeptide Myomixer is essential for satellite cell fusion and muscle regeneration. Proceedings of the National Academy of Sciences (2018).

18. Zhang, Q., et al. The microprotein Minion controls cell fusion and muscle formation. Nature Communications 8, 15664 (2017).

19. Im, W.B., et al. Differential Expression of Dystrophin Isoforms in Strains of mdx Mice with Different Mutations. Hum Mol Genet 5, 1149–1153 (1996).

20. Beastrom, N., et al. mdx5cv Mice Manifest More Severe Muscle Dysfunction and Diaphragm Force Deficits than Do mdx Mice. The American Journal of Pathology 179, 2464–2474 (2011).

21. Berger, J., Berger, S., Hall, T.E., Lieschke, G.J. & Currie, P.D. Dystrophin-deficient zebrafish feature aspects of the Duchenne muscular dystrophy pathology. Neuromuscular Disorders 20, 826–832 (2010).

22. Li, M., Hromowyk, K.J., Amacher, S.L. & Currie, P.D. Chapter 14 - Muscular dystrophy modeling in zebrafish. in Methods in Cell Biology, Vol. 138 (eds. Detrich, H.W., Westerfield, M. & Zon, L.I.) 347–380 (Academic Press, 2017).

23. Berger, J. & Currie, P.D. Zebrafish models flex their muscles to shed light on muscular dystrophies. Disease Models & Mechanisms 5, 726–732 (2012).

24. Bassett, D. & Currie, P.D. IDENTIFICATION OF A ZEBRAFISH MODEL OF MUSCULAR DYSTROPHY. Clinical and Experimental Pharmacology and Physiology 31, 537–540 (2004).

25. Bassett, D.I., et al. Dystrophin is required for the formation of stable muscle attachments in the zebrafish embryo. Development 130, 5851–5860 (2003).

26. Jin, H., et al. The dystrotelin, dystrophin and dystrobrevin superfamily: new paralogues and old isoforms. BMC Genomics 8, 19 (2007).

27. Kawahara, G., et al. Drug screening in a zebrafish model of Duchenne muscular dystrophy. Proceedings of the National Academy of Sciences 108, 5331–5336 (2011).

28. Guyon, J.R., et al. The dystrophin associated protein complex in zebrafish. Hum Mol Genet 12, 601–615 (2003).

29. Berger, J., Sztal, T. & Currie, P.D. Quantification of birefringence readily measures the level of muscle damage in zebrafish. Biochemical and Biophysical Research Communications 423, 785–788 (2012).

30. Blau, H.M., Webster, C. & Pavlath, G.K. Defective myoblasts identified in Duchenne muscular dystrophy. Proceedings of the National Academy of Sciences 80, 4856–4860 (1983).

31. Blau, H.M., Webster, C., Chiu, C.-P., Guttman, S. & Chandler, F. Differentiation properties of pure populations of human dystrophic muscle cells. Experimental Cell Research 144, 495–503 (1983).

32. Chrousos, G.P. Chapter 99 - Glucocorticoid Action: Physiology. in Endocrinology: Adult and Pediatric (Seventh Edition) (eds. Jameson, J.L., et al.) 1727-1740.e1725 (W.B. Saunders, Philadelphia, 2016).

33. Dunn, J.F. & Zaim-Wadghiri, Y. Quantitative magnetic resonance imaging of the mdx mouse model of Duchenne muscular dystrophy. Muscle & Nerve 22, 1367–1371 (1999).

34. Pratt, S.J.P., Xu, S., Mullins, R.J. & Lovering, R.M. Temporal changes in magnetic resonance imaging in the mdx mouse. BMC Research Notes 6, 262 (2013).

35. Bostock, E.L., et al. Impaired Glucose Tolerance in Adults with Duchenne and Becker Muscular Dystrophy. Nutrients 10, 1947 (2018).

36. Brazeau, G.A., Mathew, M. & Entrikin, R.K. Serum and organ indices of the mdx dystrophic mouse. Research Communications in Chemical Pathology and Pharmacology 77, 179–189 (1992).

37. Stapleton, D.I., et al. Dysfunctional Muscle and Liver Glycogen Metabolism in mdx Dystrophic Mice. PLOS ONE 9, e91514 (2014).

38. Ahmad, N., et al. Use of imaging biomarkers to assess perfusion and glucose metabolism in the skeletal muscle of dystrophic mice. BMC Musculoskeletal Disorders 12, 127 (2011).

39. Mokhtarian, A. & Even, P.C. Effect of intraperitoneal injection of glucose on glucose oxidation and energy expenditure in the mdx mouse model of Duchenne muscular dystrophy. Pflügers Archiv 432, 379 (1996).

40. Even, P.C., Decrouy, A. & Chinet, A. Defective regulation of energy metabolism in mdx mouse skeletal muscles. Biochemical Journal 304, 649–654 (1994).

41. Kaipa, B.R., et al. Dock mediates Scar- and WASp-dependent actin polymerization through interaction with cell adhesion molecules in founder cells and fusion-competent myoblasts. Journal of Cell Science 126, 360–372 (2013).

42. Namekata, K., et al. Dock3 induces axonal outgrowth by stimulating membrane recruitment of the WAVE complex. Proceedings of the National Academy of Sciences 107, 7586–7591 (2010).

43. Yajima, H. & Kawakami, K. Low Six4 and Six5 gene dosage improves dystrophic phenotype and prolongs life span of mdx mice. Development, Growth & Differentiation 58, 546–561 (2016).

44. Heller, H., Gredinger, E. & Bengal, E. Rac1 Inhibits Myogenic Differentiation by Preventing the Complete Withdrawal of Myoblasts from the Cell Cycle. Journal of Biological Chemistry 276, 37307–37316 (2001).

45. Chockalingam, P.S., et al. Dystrophin-glycoprotein complex and Ras and Rho GTPase signaling are altered in muscle atrophy. American Journal of Physiology - Cell Physiology 283, C500–C511 (2002).

46. Bai, Y., et al. Balanced Rac1 activity controls formation and maintenance of neuromuscular acetylcholine receptor clusters. Journal of Cell Science 131, jcs215251 (2018).

47. H., R.S., et al. Rac1 muscle knockout exacerbates the detrimental effect of high-fat diet on insulin-stimulated muscle glucose uptake independently of Akt. The Journal of Physiology 0(2018).

48. Samson, T., et al. Def-6, a Guanine Nucleotide Exchange Factor for Rac1, Interacts with the Skeletal Muscle Integrin Chain α7A and Influences Myoblast Differentiation. Journal of Biological Chemistry 282, 15730–15742 (2007).

49. Vieira, Natassia M., et al. Jagged 1 Rescues the Duchenne Muscular Dystrophy Phenotype. Cell 163, 1204–1213 (2015).

50. Thisse, C. & Thisse, B. High-resolution in situ hybridization to whole-mount zebrafish embryos. Nat. Protocols 3, 59–69 (2008).

51. Wildforster, V. & Dekomien, G. Detecting copy number variations in autosomal recessive limb-girdle muscular dystrophies using a multiplex ligation-dependent probe amplification (MLPA) assay. Mol Cell Probes 23, 55 – 59 (2009).

52. Hightower, R.M., et al. The SINE Compound KPT-350 Blocks Dystrophic Pathologies in DMD Zebrafish and Mice. Molecular Therapy (2019).

53. Schindelin, J., et al. Fiji: an open-source platform for biological-image analysis. Nature Methods 9, 676–682 (2012).

54. Ahrens, H.E., et al. Analyzing Satellite Cell Function During Skeletal Muscle Regeneration by Cardiotoxin Injury and Injection of Self-delivering siRNA In Vivo. JoVE, e60194 (2019).

55. Koster, J. & Rahmann, S. Snakemake--a scalable bioinformatics workflow engine. Bioinformatics 28, 2520–2522 (2012).

56. Patro, R., Duggal, G., Love, M.I., Irizarry, R.A. & Kingsford, C. Salmon provides fast and bias-aware quantification of transcript expression. Nat Methods 14, 417–419 (2017).

57. Ewels, P., Magnusson, M., Lundin, S. & Kaller, M. MultiQC: summarize analysis results for multiple tools and samples in a single report. Bioinformatics 32, 3047–3048 (2016).

58. Soneson, C., Love, M.I. & Robinson, M.D. Differential analyses for RNA-seq: transcript-level estimates improve gene-level inferences. F1000Res 4, 1521 (2015).

59. Love, M.I., Huber, W. & Anders, S. Moderated estimation of fold change and dispersion for RNA-seq data with DESeq2. Genome biology 15, 550 (2014).

60. Livak, K.J. & Schmittgen, T.D. Analysis of Relative Gene Expression Data Using Real-Time Quantitative PCR and the 2-[Delta][Delta]CT Method. Methods 25, 402–408 (2001).

61. Cheung, T.H., et al. Maintenance of muscle stem-cell quiescence by microRNA-489. Nature 482, 524–528 (2012).

62. Gharaibeh, B., et al. Isolation of a slowly adhering cell fraction containing stem cells from murine skeletal muscle by the preplate technique. Nat. Protocols 3, 1501–1509 (2008).

63. Alexander, M.S., et al. MicroRNA-199a is induced in dystrophic muscle and affects WNT signaling, cell proliferation, and myogenic differentiation. Cell Death Differ 20, 1194–1208 (2013).

64. Leikina, E., et al. Myomaker and Myomerger Work Independently to Control Distinct Steps of Membrane Remodeling during Myoblast Fusion. Developmental Cell 46, 767-780.e767 (2018).

65. Vadivelu, S.K., et al. Muscle Regeneration and Myogenic Differentiation Defects in Mice Lacking TIS7. Molecular and Cellular Biology 24, 3514–3525 (2004).

66. Sievers, F. & Higgins, D.G. Clustal Omega for making accurate alignments of many protein sequences. Protein Science 27, 135–145 (2018).

