## Supplementary figures and images for "“DOCK3 is a dosage-sensitive regulator of skeletal muscle and Duchenne muscular dystrophy-associated pathologies”"

### Figure 3

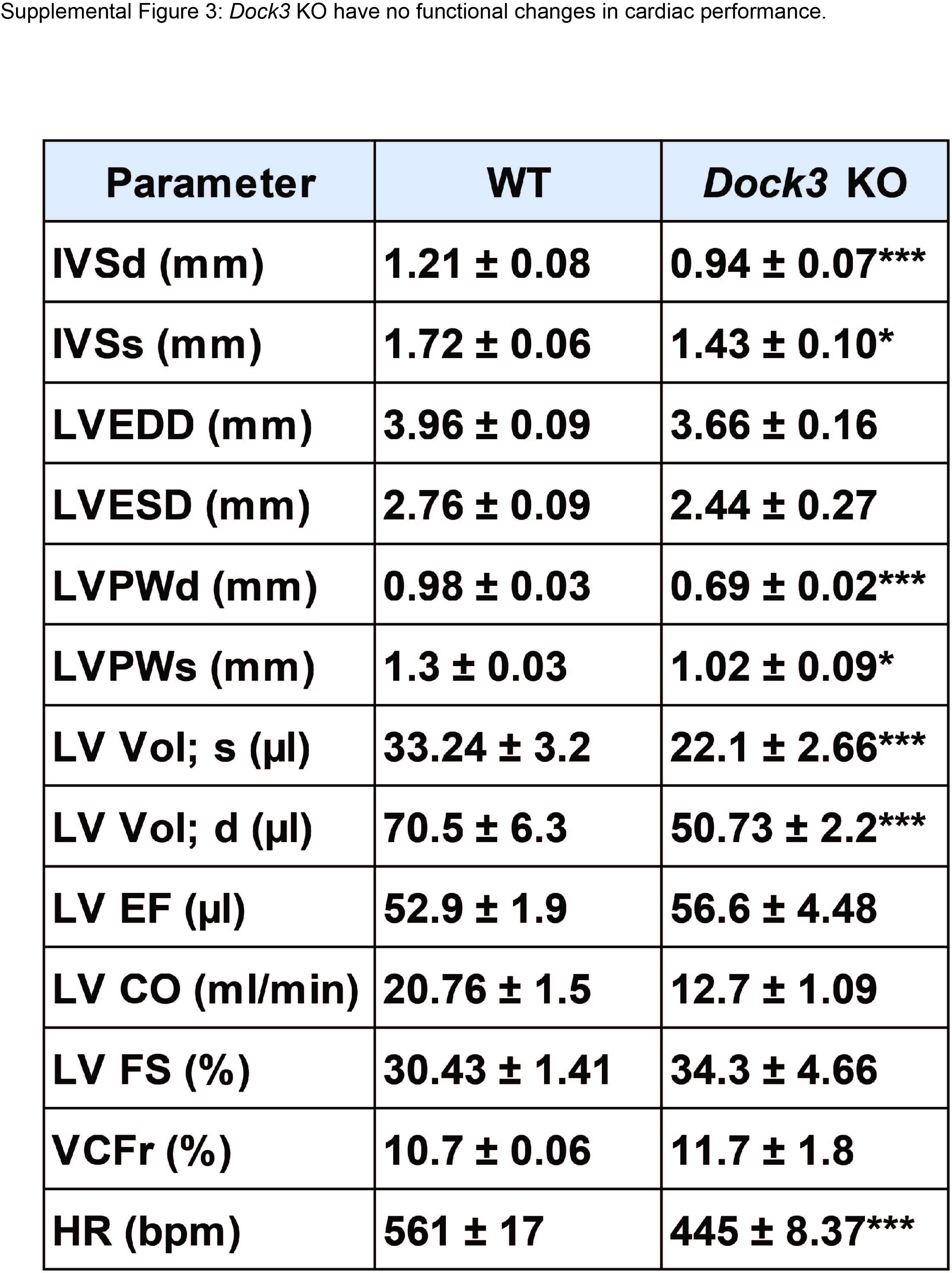

### Supplemental Figure 1

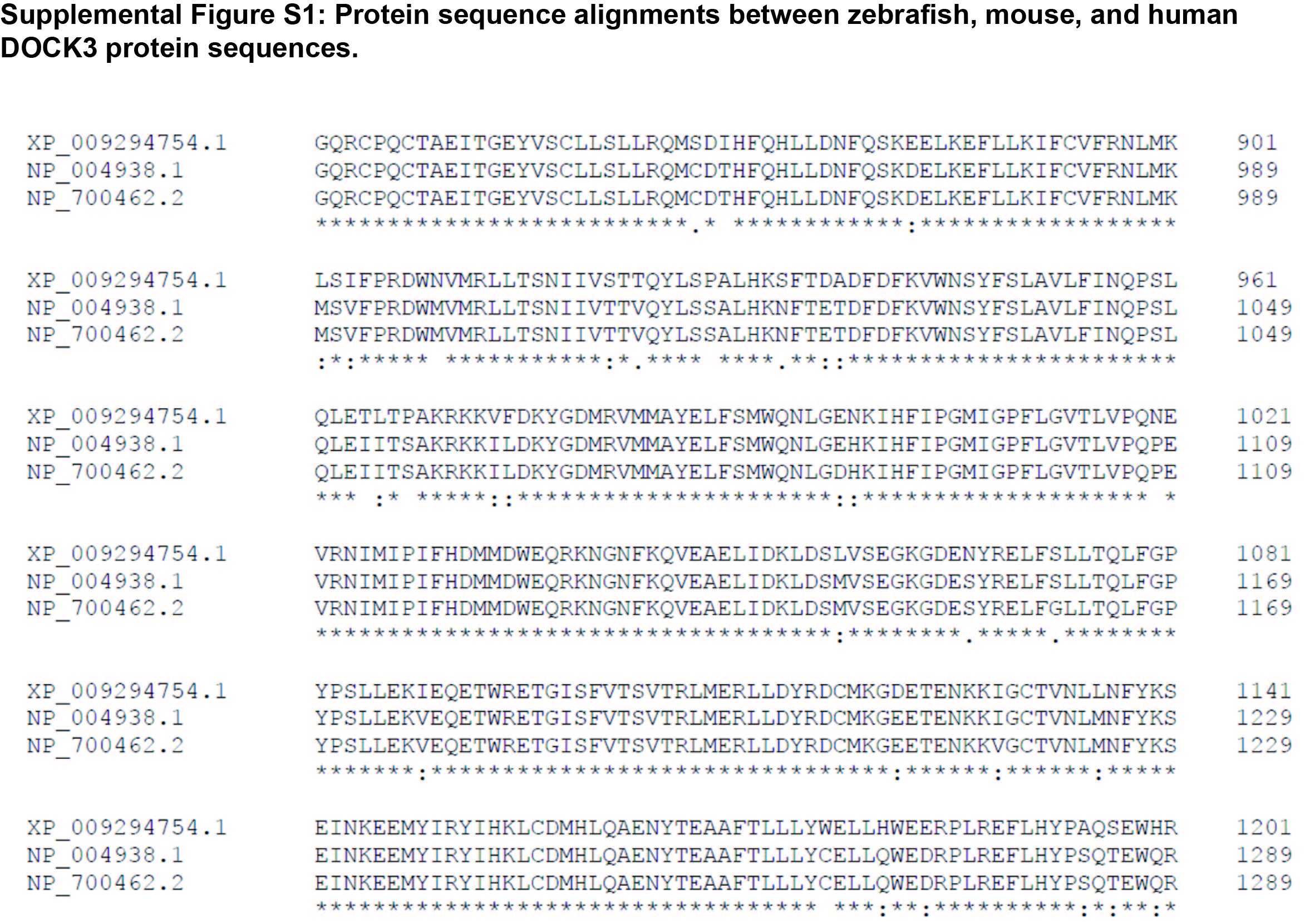

### Supplemental Figure 2

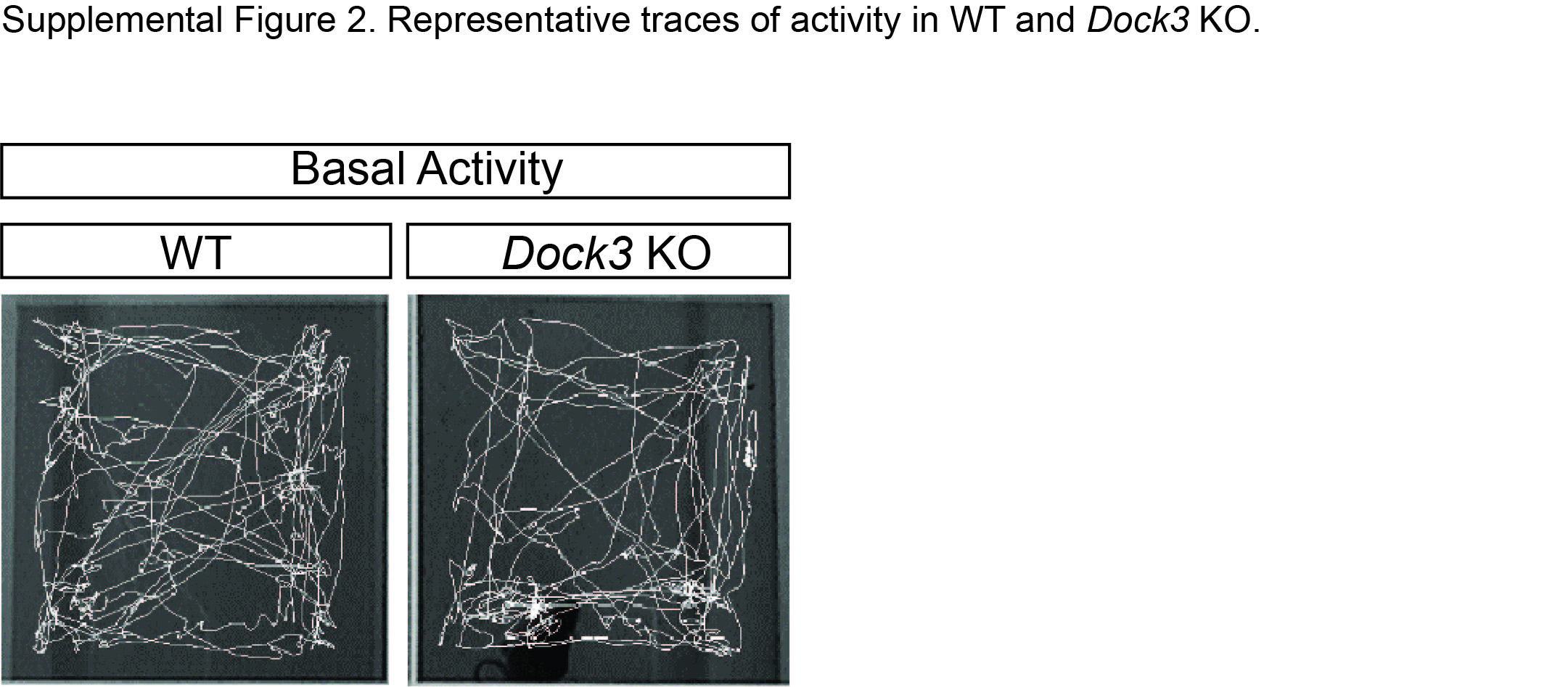
